# Bioinspired Bimetallic Ions Functionalized MOF SAzyme Nanocomposites for Synergistic Ferroptosis/Cuproptosis-Enhanced Immune Checkpoint Therapy

**DOI:** 10.1101/2024.08.12.607118

**Authors:** Linjiao Yang, Mengmeng Pan, Haofan Hu, Furong Liu, Ming Jiang, Shangwu Ning, Xiaoping Chen, Zhanguo Zhang, Xu Yu, Li Xu

## Abstract

Immune checkpoint blockade (ICB) generates sustained responses in immunogenic cancers, but its effectiveness is limited in tumors lacking immune activity. Here, we construct a bioinspired bimetallic ions functionalized nanoscale metal-organic framework (NMOF) single-atom nanozyme (SAzyme) loaded with doxorubicin (Dox) (NMOF-Fe/Cu-Dox nanocomposite) to effectively trigger anti-tumor immune responses while addressing the immunosuppressive tumor microenvironment (TME). The NMOF-Fe/Cu-Dox nanocomposite has been demonstrated to efficiently reverse the TME by generating reactive oxygen species and oxidizing glutathione. Camouflaging NMOF-Fe/Cu-Dox nanocomposites with bioinspired cancer cell membrane (NMOF-Fe/Cu-Dox@M) enables its navigation to the tumor region through homologous targeting. The highly efficient uptaken by cancer cells selectively induced synergistic ferroptosis and cuproptosis in these cells. Furthermore, *in vitro* and *in vivo* experiments demonstrate that the ferroptosis in cancer cells can polarize tumor-associated macrophages (TAMs) towards anti-tumoral M1 phenotype and significantly diminish pro-tumoral M2 phenotype. We find that NMOF-Fe/Cu-Dox@M could induce the ferroptosis of M2 macrophages, while no effect to M1 macrophages. In addition, a significant increase of anti-tumor infiltrating CD8^+^ T cells, while a remarkable decrease of CD4^+^ regulatory T cells were observed. These findings suggested that NMOF-Fe/Cu-Dox@M could efficiently modulate TME, enhance tumoricidal immunity and elevate the therapeutic efficiency of ICB. Moreover, the combination of NMOF-Fe/Cu-Dox@M with αPD-1 effectively eradicated hepatocellular carcinoma cells *in vivo*, outperforming the use of either NMOF-Fe/Cu-Dox@M or αPD-1 alone. In summary, our study presents a therapeutic strategy that leverages coordinated ferroptosis and cuproptosis with therapeutic efficiency of ICB, underscoring the promise of combined chemoimmunotherapy.

## Introduction

Cancer immune checkpoint blockade (ICB) therapy harnesses the capabilities of the host immune system to provoke extensive anti-tumor reactions and establish long-term immune surveillance^1–3^. Typically, the ICB that targets T-lymphocyte immunosuppressive pathways has shown promising response rates with minimal side effects in various cancers^4–6^. However, it has not yet been proven to provide survival benefits in lowly immunogenic tumors^7, 8^, such as hepatocellular carcinoma (HCC) and pancreatic cancers^9, 10^. Meanwhile, immunosuppressive tumor microenvironment (TME), characterized by overexpressed tumor molecules like CD47 and PD-L1 in tumor cells, hinders the activation of the immune system and leads to low response to ICB^4, 11–13^. Therefore, it is imperative to create novel clinically applicable immunogenic treatment modalities that can serve as immunostimulatory adjuvants to enhance the effectiveness of ICB.

Currently, inducing apoptosis by some anti-cancer drugs or nano-medications has emerged as a primary approach for cancer therapy^14, 15^. However, apoptosis is commonly recognized as a type of immune-tolerant cell death, since the removal of apoptotic tumor cells by macrophages may enhance the immunosuppression and hinder the activation of the host immune system^16, 17^. Thus, triggering alternative non-apoptotic cell death pathways, including necroptosis^18^, pyroptosis^19^, ferroptosis and cuproptosis^20–22^, represents a novel approach to eliminate tumor cells, bolster tumor immunogenicity^17, 23^, and consequently stimulate the anti-tumor immunogenic efficacy. Among them, ferroptosis and cuproptosis have attracted significant attention due to their high efficiency in inducing programmed cell death^20, 24^. Additionally, the immunogenic treatments via ferroptosis and cuproptosis pathways have the potential to serve as immunostimulatory adjuvants to enhance the efficacy of ICB, but the relevant anti-tumor activity and mechanism were not investigated.

Ferroptosis is a new form of regulated cell death dependent on iron ion dyschondrosteosis and excessive reactive oxygen species (ROS), characterized by inducing lipid peroxidation (LPO), and morphologically-shrunken mitochondria with reduced or disappeared cristae^25–27^. In contrast, cuproptosis is programmed to be driven by the accumulation of intracellular copper ions (Cu^2+^)^28–31^. The upstream regulator, mitochondrial ferredoxin 1 (FDX1), reduces divalent Cu^2+^ to monovalent copper ions (Cu^+^), which then bind to dihydrolipoamide S-acetyltransferase (DLAT) and interfere the Fe-S cluster proteins. Additionally, along with intermediates in the lipoic acid pathway, Cu^+^ facilitates the lipoylation of enzymes containing the pyruvate dehydrogenase complex in the tricarboxylic acid (TCA) cycle, leading to proteotoxic stress and cell death^31^. Although ferroptosis and cuproptosis were initiated by different triggers and mechanisms, they shared some commonalities in anti-tumor therapy, such as oxidative stress, abnormal metabolism, perturbation of cellular processes and mitochondrial damage. These effects could be particularly significant since disruption of metal ion homeostasis and the redox balance is supposed to result in cellular damage and cell death^20, 24^. Thus, the combination of ferroptosis and cuproptosis might exert the better anticancer effect than either cuproptosis or ferroptosis alone^20, 24, 32^. However, whether and how ferroptosis and cuproptosis synergistically induce immunogenic cell death, modulate the TME, affect macrophage polarization and activation, and activate T cell proliferation and infiltration, thereby enhancing the efficacy of ICB, remains unknown and requires further exploration.

In recent years, single-atom nanoenzyme (SAzyme) with highly catalytic activity has garnered widespread interests among researchers^33–35^. In these SAzymes, the uniformly dispersed metal atom active centers could maximize the utilization efficiency of atoms and enhanc the enzymatical catalytic activity. With the increasing interests, some SAzymes have been exploited for the organic catalysis synthesis^36, 37^, biosensors construction^38^, antibacterial treatment and cancer therapy^26, 39^. Despite significant efforts in this area, the challenge of rapid, easy, and efficient construction of SAzymes with synergistic ferroptosis and cuproptosis simutaneously to induce programmed cancer cell death, modulate the TME, and activate immunotherapy, particularly to enhance ICB, remains a major obstacle. We recently reported a new class of metallic ions functionlized nanosized metal organic framerwork (NMOF) nanozymes, ranging from single metal ions to bimetallic ions functionlization, for successful antibacterial and photodynamic therapy (PDT) of cancers^40, 41^. Upon deeper investigation, we found these metallic ions functionalized NMOFs might belong to SAzymes. The NMOF exhibits significantly higher chelating efficiency for various metal ions, leading to the creation of NMOF-M^n+^ SAzymes with excellent uniform atomic dispersion and superior enzyme-like activities compared to other SAzymes. By chelating *mono*-, *bi*-, or *tri*-metallic ions, we can tailor the multi-enzyme activities of the SAzymes, while the mesoporous structure of the NMOF facilitates the loading of anti-tumor drugs for synergistic cancer treatment. Inspiriated by these findings, we were motivated to develop a Fe^3+^ and Cu^2+^ bimetallic ions functionlized NMOF SAzyme, designated as NMOF-Fe/Cu SAzyme, to incorporate the ferroptosis and cuproptosis, for the catalytic therapy of HCC. We hypothesized that the synergistic effects of ferroptosis and cuproptosis might trigger a stronger local immune response, while cooperating with ICB to achieve systemic and effective HCC therapy.

Herein, we further designed nanocomposites based on NMOF-Fe/Cu SAzyme with the anticancer drug doxorubicin (Dox) loading and bioinspired HCC cell membrane coating, denoted as NMOF-Fe/Cu-Dox@M, for the HCC treatment (Scheme 1). In this design, the NMOF-Fe/Cu SAzyme is expected to display peroxidase (POD) and glutathione peroxidase (GPx) enzyme-like activities, which could efficiently generate the ROS and deplete the GSH in the tumor cells. Thus, the excessive ROS could induce the LPO and the reduced Cu^+^ accumulation was capable of inducing ferroptosis and cuproptosis, while incorporating Dox into NMOF-Fe/Cu SAzyme to enhance therapeutic efficacy. We employed bioinspired cancer cell membranes to encapsulate the NMOF-Fe/Cu-Dox, thereby achieving homologous targeting towards HCC tumor cells and prolonging the circulattion time of the nanocomposites *in vivo*^42–44^. Interesting, we found that synergistic ferroptosis and cuproptosis in cancer cells could polarize tumor-associated macrophages (TAMs) towards anti-tumoral M1 macrophages and significantly diminish pro-tumoral M2 macrophages, while having no effect on M1 macrophages. Additionally, there was a significant increase in anti-tumor infiltrating CD8^+^ T cells and a remarkable decrease in CD4^+^ regulatory T cells, demonstrating that NMOF-Fe/Cu-Dox@M successfully activated the anti-tumor immunity. Furthermore, the mechanisms by which synergistic ferroptosis and cuproptosis in cancer cells effectively reversed immunosuppressive TME, activated immune response, and enhanced ICB efficacy were demonstrated.

**Scheme 1.**
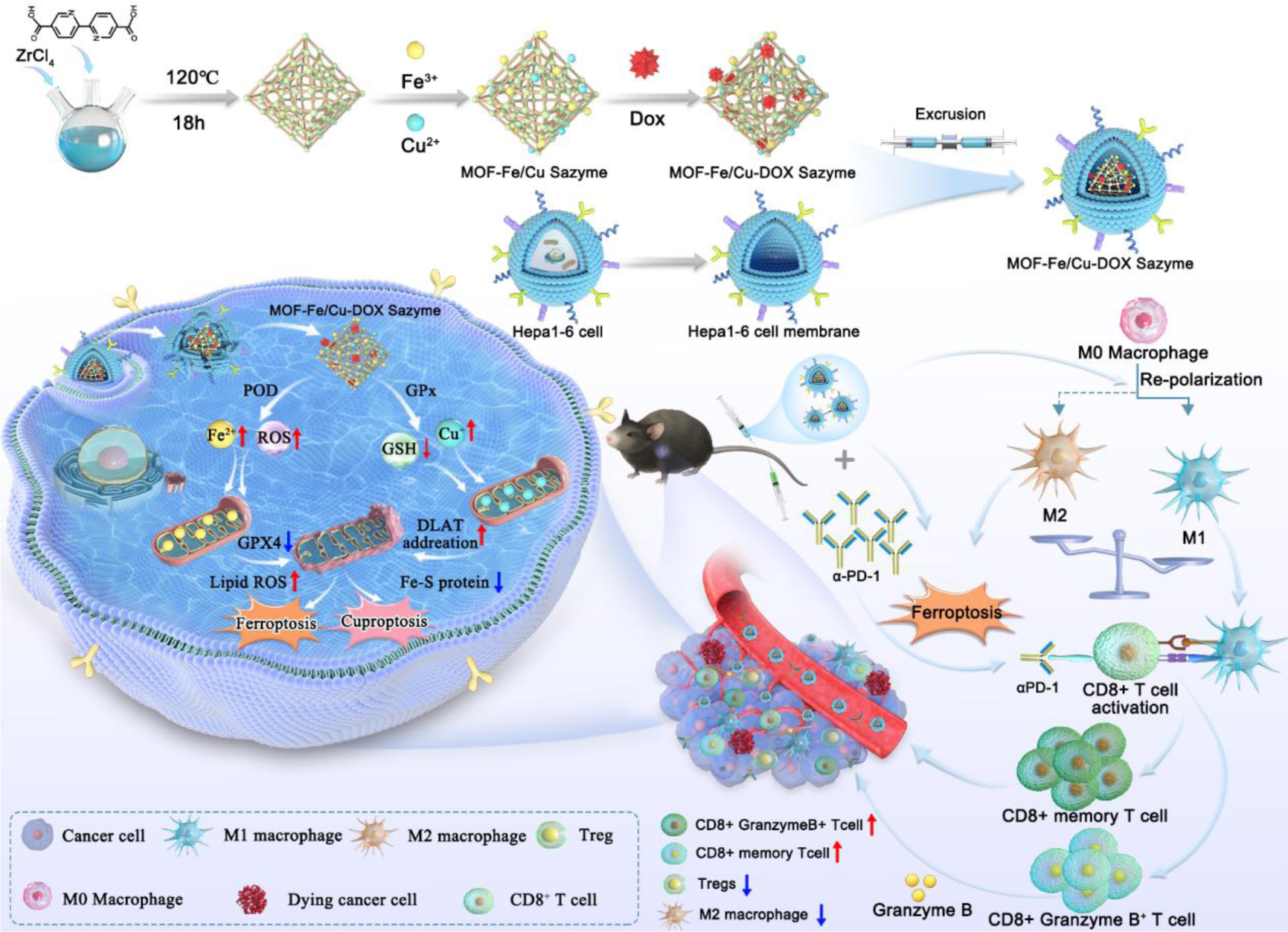
Schematic illustration of synthesis of the NMOF-Fe/Cu-Dox@M and the synergistic therapeutic mechanism to HCC.

## Results

### Construction of Fe/Cu bimetallic ions functionalized NMOF single-atom nanozyme (NMOF-Fe/Cu SAzyme)

To construct the NMOF-Fe/Cu SAzyme, the 2,2’-bipyridine-5,5’-dicarboxylic acid was chosen as the building block because of the highly chelating capacity of the bipyridine group with the metal ions^40, 41^. The NMOF, i.e., nano-sized UIO-67 NMOF nanoparticles, was synthesized via a one-step hydrothermal reaction (Zr^4+^ and ligand), as described in **Scheme 1**, exhibiting an octahedral three-dimensional structure (Fig. 1a) with the average particle size of ∼200 nm (Fig. 1b). Subsequently, the bimetallic ions functionalized NMOF nanozyme was obtained by chelating the Fe^3+^ and Cu^2+^ ions with bipyridine ligands on the NMOF. SEM images revealed that the NMOF-Fe/Cu retained the UIO-67 morphology after chelating the Fe^3+^/Cu^2+^ ions (Supplementary Fig. S1a). The high-angle annular dark-field scanning transmission electron microscopic (HAADF-STEM) images demonstrated the uniform distribution of Zr, Cu, and Fe on the NMOF-Fe/Cu (Fig. 1c). The aberration-corrected elemental mapping image clearly showed that Fe and Cu were atomically dispersed on NMOF-Fe/Cu (marked by red circles), which could be assigned to Fe and Cu single atoms (Fig. 1d). The contents of optimized Fe^3+^ and Cu^2+^ ions on the NMOF-Fe/Cu were determined to be 0.022 and 1.21 μmol/mg by inductively coupled plasma mass spectrometry (ICP-MS, Supplementary Table S1). Thermogravimetric analysis revealed a reduced weight loss for NMOF-Fe/Cu compared to NMOF, confirming the successful chelation of Fe and Cu on the nanozyme (Supplementary Fig. S3). Fourier transform infrared (FTIR) spectroscopy was also employed to disclose the functional groups on the NMOF and NMOF-Fe/Cu (Supplementary Fig. S4). The C=N stretching vibrations at 1596 cm^-1^, C-N stretching vibrations at 1422 cm^-1^, the μ_3_-O bonds stretching at 648 cm^-1^ and C-H absorption at 775 cm^-1^ were observed on the NMOF and NMOF-Fe/Cu SAzyme^41^.

**Fig 1.**
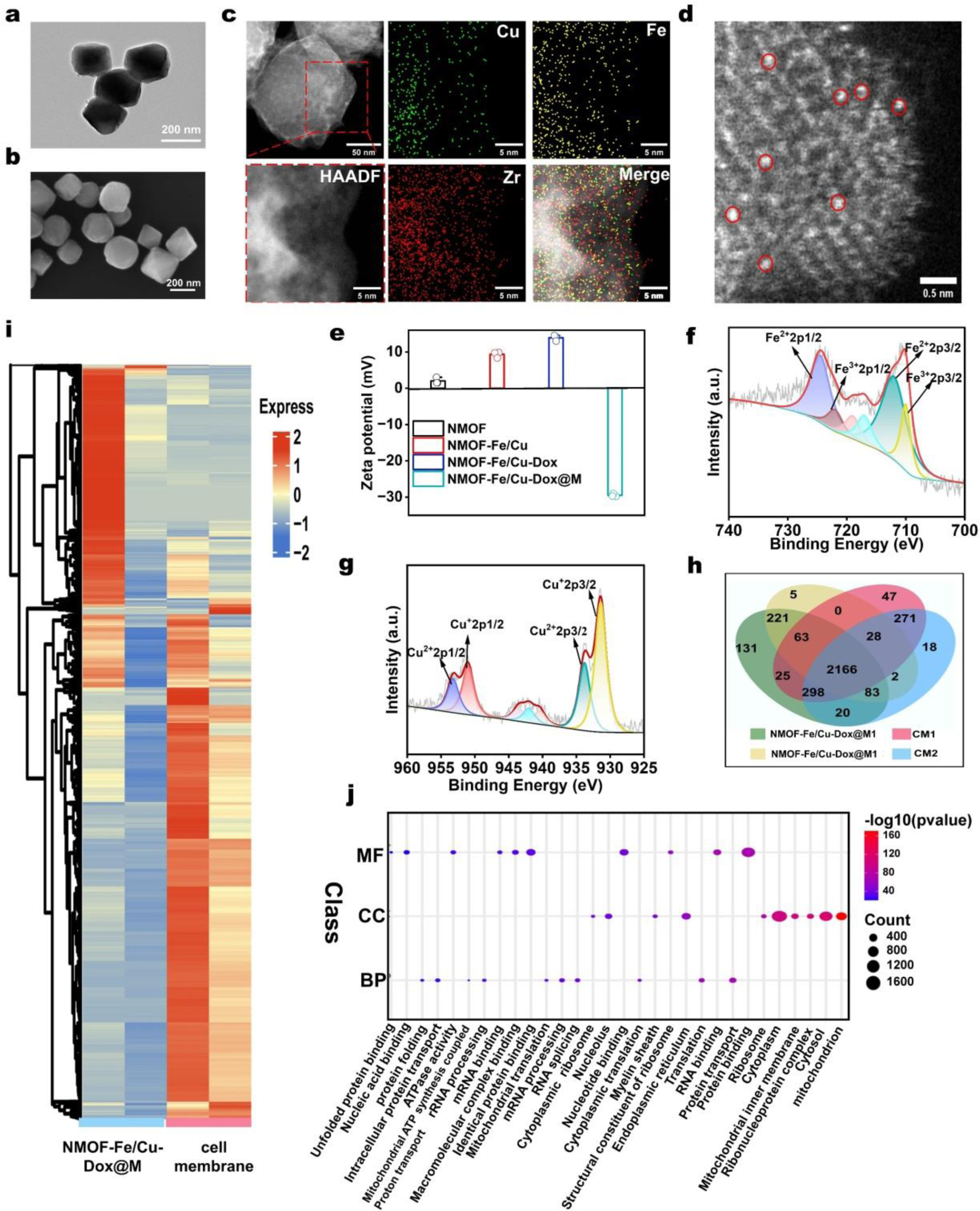
Characterization of the NMOF-Fe/Cu-Dox@M. **a**, **b**, The TEM and SEM images of NMOF. **c**, HAADF-STEM images and element mapping of the NMOF-Fe/Cu SAzyme. **d**, Fe and Cu dispersed on NMOF-Fe/Cu SAzyme (marked by red circles). **e**, ζ-potentials of NMOF, NMOF-Fe/Cu, NMOF-Fe/Cu-Dox and NMOF-Fe/Cu-Dox@M. **f**, **g**, XPS analysis of Fe 2p and Cu 2p of NMOF-Fe/Cu SAzyme. **h**, Venn diagram showing the overlap of 2166 proteins in Hepa1-6 cells membrane and NMOF-Fe/Cu-Dox@M. **i**, Heatmap of proteins in Hepa1-6 cells membrane and NMOF-Fe/Cu-Dox@M. **j**, GO enrichment analysis of proteins in NMOF-Fe/Cu-Dox@M. The top 10 processes were displayed.

X-ray photoelectron spectroscopy (XPS) survey was used to investigate the chemical environment of NMOF-Fe/Cu SAzyme. The full XPS data showed characteristic peaks of Zr 3d, O 1s, N 1s, and C 1s in NMOF and NMOF-Fe/Cu (Supplementary Fig. S5). The extra Cu (931 eV) and Fe (709 eV) regions appeared on the NMOF-Fe/Cu nanozyme (Supplementary Fig. S5). Further analysis of the Fe 2p and Cu 2p in the high-resolution XPS spectra revealed the presence of multivalent Fe (+2 and +3) and Cu (+1 and +2) species on the NMOF-Fe/Cu SAzyme (Fig. 1f and Fig. 1g). The binding energies could be attributed to the Fe^2+^ (2p^3/2^, 712.0 eV and 2p^1/2^, 724.5 eV), Fe^3+^ (2p^3/2^, 710.0 eV and 2p^1/2^, 722.0 eV) and Cu^+^ (2p^3/2^, 933.8 eV and 2p^1/2^ 953.2 eV) and Cu^2+^ (2p^3/2^, 931.4 and 2p^1/2^ 951.2 eV), respectively. In contrast, the Zr 3p and O 1s XPS peaks of NMOF remained unchanged, suggesting no coordination between the Zr-O cluster and Fe/Cu ions. In addition, the X-ray diffraction (XRD) patterns illustrated that there were no crystal changes (Supplementary Fig. S6). In summary, these findings underscored the successful construction of the Fe^3+^/Cu^2+^ bimetallic ions-functionalized NMOF SAzyme.

### Construction of NMOF-Fe/Cu-Dox@M

For the HCC therapy, the Dox drug was loaded into the mesoporous pores on the NMOF-Fe/Cu SAzyme for synergistic therapy (termed as NMOF-Fe/Cu-Dox, Supplementary Fig. S1b). To enhance the biocompatibility, cellular immune escape and homologous targeted cancer ability, we adopted cancer cell membrane to coat the nanoparticles via one-step extrusion method (Supplementary Fig. S2). The prepared cancer cell membrane encapsulating NMOF-Fe/Cu-Dox nanoparticles (NMOF-Fe/Cu-Dox@M) were measured through dynamic light scattering (DLS) for ζ-potential and particle size analysis. The observed increase in particle sizes and alterations in ζ-potentials provided evidence for the successful construction of NMOF-Fe/Cu-Dox@M (Fig. 1e, Supplementary Fig. S7). As shown in Supplementary Fig. S8, the ζ-potentials and particle sizes of NMOF-Fe/Cu-Dox@M maintained their stability for 7 days in water and DMEM medium (including 10% FBS), proving that NMOF-Fe/Cu-Dox@M was with ignorable degradation. Meanwhile, previous studies have demonstrated that some relative proteins, including CD47 and E-cadherin, present in cancer cell membranes played a crucial role in mediating cellular immune escape and homologous targeted effects^42, 45^. Thus, we used the sodium dodecyl sulfate polyacrylamide gel electrophoresis (SDS-PAGE) and proteomic analysis to investigate the protein composition and differences between the cancer cell and NMOF-Fe/Cu-Dox@M. In Supplementary Fig. S9, we observed almost identical protein gel bands in both Hepa1-6 cell membrane and NMOF-Fe/Cu-Dox@M, suggesting the preservation of membrane proteins on the nanoparticles. Then, the proteomic results revealed that a total of 3393 proteins were detected in the nanoparticles and Hepa1-6 cell membranes (Fig. 1h). Among these proteins, 2166 were found to be overlapped in both groups. Considering the sequence coverage being > 50%, the 325 proteins of the overlapped ones were considered to be primarily reserved in NMOF-Fe/Cu-Dox@M, and the expressions of these proteins are illustrated in Fig. 1i. We then utilized gene ontology (GO) functional enrichment analysis to explore the distinct biological and molecular functions associated with these proteins. The results highlighted a significant involvement of these proteins in biological processes related to protein transport, as well as molecular functions such as protein binding and nucleic acid binding (Fig. 1j). These outcomes collectively suggested the successful preservation of cancer cell membrane proteins on the NMOF-Fe/Cu-Dox@M, with a substantial number actively participating in homologous targeted and adhesive processes.

### Catalytic activity of NMOF-Fe/Cu SAzyme

In previous studies, we found that the metal ions functionalized NMOF nanozyme showed the multiple enzyme-mimic property^40, 41^. Herein, we investigated the enzyme catalytic activity of the NMOF-Fe/Cu SAzyme. Initially, we evaluated the POD and OXD-mimic activities with the TMB reaction. As shown in Fig. 2a, the NMOF-Fe/Cu SAzyme exhibited catalytical oxidization of the TMB in the presence of H_2_O_2_, resulting in the production of the blue oxidized TMB product, TMB_ox_. In contrast, no colored TMB_ox_ was formed in the absence of H_2_O_2_. These observations underscored that the NMOF-Fe/Cu SAzyme possessed POD-mimic enzymatic activity, but not the OXD-mimic activity. Subsequently, we used the oxidation of methylene blue (MB) and terephthalic acid (TA) to verify the generation of •OH in the system. As depicted in Fig. 2b, treatment with either H_2_O_2_ or NMOF-Fe/Cu SAzyme in MB reaction system resulted in almost no change in the absorption intensity of MB compared to that of the original MB solution. However, it was sharply weakened in the coexistence of MB, H_2_O_2_, and NMOF-Fe/Cu SAzyme, which revealed the generation of •OH and oxidation of MB, and the occurrence of Fenton reaction. In addition, the oxidation of TA to form the fluorescent product, 2-hydroxy terephthalic acid (TA_OH_) was observed (Supplementary Fig. S10), further supporting the generation of •OH by the NMOF-Fe/Cu SAzyme and H_2_O_2_. Also, the generation of •OH was verified by the electron spin resonance (ESR) spectra. The intermediate radical-DMPO adducts exhibited the 1:2:2:1 characteristic signal^46^, suggesting the production of •OH (Fig. 2c). The above results demonstrated that the as-prepared NMOF-Fe/Cu SAzyme possessed intrinsic Fenton-like activity and could effectively catalyze the decomposition of H_2_O_2_ into highly toxic •OH, thus exerting potential for tumor treatment.

**Fig 2.**
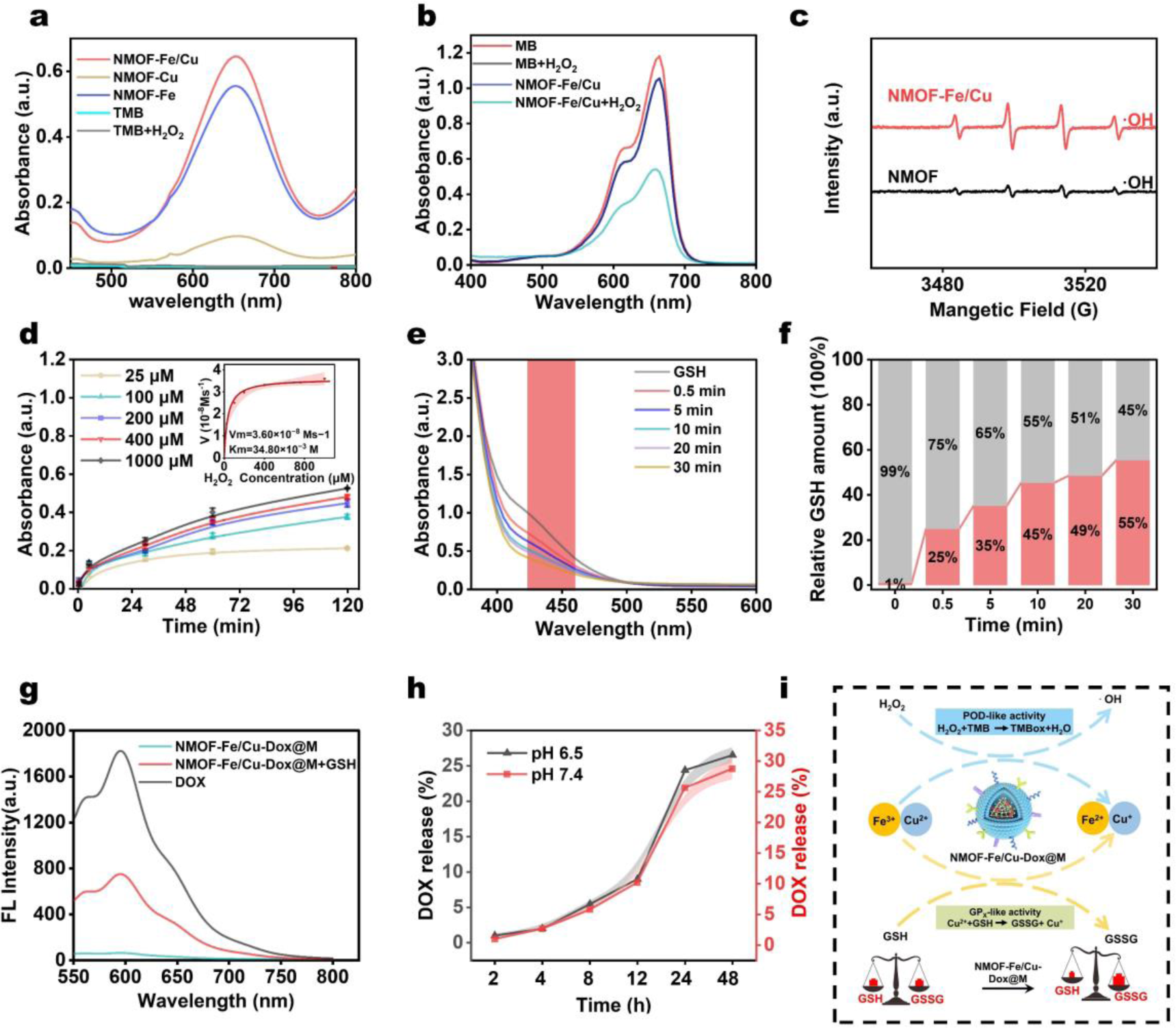
Evaluation of the enzyme-like catalytic activity of NMOF-Fe/Cu SAzyme and the drug release from the NMOF-Fe/Cu-Dox nanocomposites. **a**, The oxidation of the TMB by NMOF-Fe, NMOF-Cu and NMOF-Fe/Cu SAzyme, respectively. **b**, Characterization of the Fenton reaction with •OH generation in the NMOF-Fe/Cu SAzyme in the absence or presence of H_2_O_2_. **c**, ESR spectra of •OH generation by NMOF-Fe/Cu SAzyme and NMOF, respectively. **d**, The reaction-time curves of TMB colorimetric reaction catalyzed by NMOF-Fe/Cu SAzyme. Right, the magnified initial linear portion of the nanozyme reaction-time curves. **e**, **f**, The depletion of GSH by NMOF-Fe/Cu SAzyme. **g**, The fluorescence spectra of Dox, NMOF-Fe/Cu-Dox and NMOF-Fe/Cu-Dox+GSH. **h**, The Dox release profile from NMOF-Fe/Cu-Dox at pH of 6.5 and 7.4 PBS and their corresponding release percent over time. **i**, The mechanism of NMOF-Fe/Cu-Dox@M with multi-enzyme catalytic activities and induction of redox homeostasis for HCC therapy.

The POD catalytic activity of the NMOF-Fe/Cu SAzyme could be modulated by chelating of different ratios of Fe^3+^ and Cu^2+^ ions (Supplementary Fig. S11). Meanwhile, the bimetallic ions NMOF-Fe/Cu SAzyme exhibited higher catalytic activity than that of NMOF-Fe or NMOF-Cu SAzyme (Fig. 2a). Notably, the optimal POD-mimic catalytic activity was achieved at a Fe^3+^/Cu^2+^ ratio of 2:1. The POD activity increased with the prolonging incubation time with NMOF-Fe/Cu SAzyme (Supplementary Fig. S12). Under the optimal condition, we examined the steady-state catalytic kinetics of NMOF-Fe/Cu SAzyme across various concentrations of H_2_O_2_. The Michaelis-Menten equation was obtained with the *K_M_* and *V_max_* values of 34.80 × 10^−3^ M and 3.70 × 10^−8^ M s^−1^, respectively (Fig. 2d), exhibiting an excellent catalytical activity of the NMOF-Fe/Cu SAzyme.

As previous studies reported, the Cu^2+^ ions can react with GSH to generate the Cu^+^ and oxidized glutathione (GSSG)^47, 48^. To determine whether NMOF-Fe/Cu SAzyme can effectively consume GSH, we analyzed the GSH content in the supernatant using 5,5’-dithiobis (2-nitrobenzoic acid) (DTNB) as a probe after filtrating out the NMOF-Fe/Cu SAzyme^41^. Our results showed that GSH levels significantly decreased over time (Fig. 2e and Fig. 2f), with 45% of 5 mM GSH being consumed by NMOF-Fe/Cu SAzyme (100 μg mL^−1^). It is also noteworthy that NMOF-Cu exhibited better GPx-like activity in consuming GSH compared to NMOF-Fe (Supplementary Fig. S13). However, the NMOF-Fe/Cu SAzyme showed a higher GPx-like activity than NMOF-Cu, even though some of the Cu were replaced by Fe ions. This indicates a synergistical enhancement of the GPx-like activity in the NMOF-Fe/Cu SAzyme. These findings demonstrate that our NMOF-Fe/Cu SAzyme had highly efficient POD- and GPx-like activities and was expected to be utilized for tumor therapy.

### Evaluation of the Dox loading and release from NMOF-Fe/Cu-Dox

To enhance the synergistic efficiency of cancer treatment, we incorporated an antitumor drug, Dox, into the NMOF-Fe/Cu SAzyme for the chemodynamic therapy. We assessed the drug loading efficiency and achieved a loading capacity of 13.3% (153 μg/mg of NMOF) (Supplementary Fig. S14). Notably, the coordination between Dox and the metal ions caused a decrease in the fluorescence intensity of Dox (Fig. 2g). To address this, we used GSH to restore the fluorescence signal of Dox, as reported in previous studies^49^. As shown in Fig. 2g, our NMOF-Fe/Cu-Dox exhibited distinct sustained release behaviors for Dox in both pH 7.4 and pH 6.5 PBS, comparable to each other. Ultimately, the overall release content of Dox reached 28% at pH 7.4 and 26% at pH 6.5 after 48 h of incubation (Fig. 2h). In summary, the proposed catalytic mechanism of NMOF-Fe/Cu-Dox@M for the generation of the POD-like activity (•OH) and GPx-like activity (depletion of GSH) was comprehensively displayed in Fig. 2i.

### Homologous tumor-targeted delivery of NMOF-Fe/Cu-Dox@M

To access the homologous targeting ability of the cancer cell membrane coated nanoparticles towards the cancer cells^42–44, 50^, murine hepatoma cancer cell line, Hepa1-6, and two normal cell lines, mouse embryonic fibroblast cells, NIH3T3, and human umbilical vein endothelial cells, HUVEC, were separately incubated with NMOF-Fe/Cu-Dox@M and NMOF-Fe/Cu-Dox, and the uptake efficiency was compared by the normal and tumor cells. As depicted in Fig. 3a-c, the results revealed that there was a substantial augmentation in the internalization of NMOF-Fe/Cu-Dox@M when coated with the bioinspired cell membrane of Hepa1-6 cells, in contrast to the control group lacking the membrane decoration. At 8 h, the uptake of NMOF-Fe/Cu-Dox@M by Hepa1-6 cells was two-fold of that of the control group (NMOF-Fe/Cu-Dox), indicating the excellent homologous targeting ability of NMOF-Fe/Cu-Dox@M towards cancer cells (*p*=0.0013). On the contrary, incubating NMOF-Fe/Cu-Dox@M with normal NIH3T3 and HUVEC cells for 1 h, 4 h, and 8 h resulted in significantly lower uptake of NMOF-Fe/Cu-Dox@M compared to the NMOF-Fe/Cu-Dox (Fig. 3d-f, Supplementary Fig. S15 and S16). After an 8 h incubation, the fluorescence intensity was approximately only half of the NMOF-Fe/Cu-Dox group. These findings underscored that the cancer cell membrane coating significantly promoted the delivery of nanoparticles to the corresponding cancer cells, while simultaneously reduced uptake by normal cells, thus decreasing the biotoxicity of the nanoparticles. According to previous proteomic results, cancer cell proteins retained in the outer membrane of NMOF-Fe/Cu-Dox@M might contribute to this outcome.

**Fig 3.**
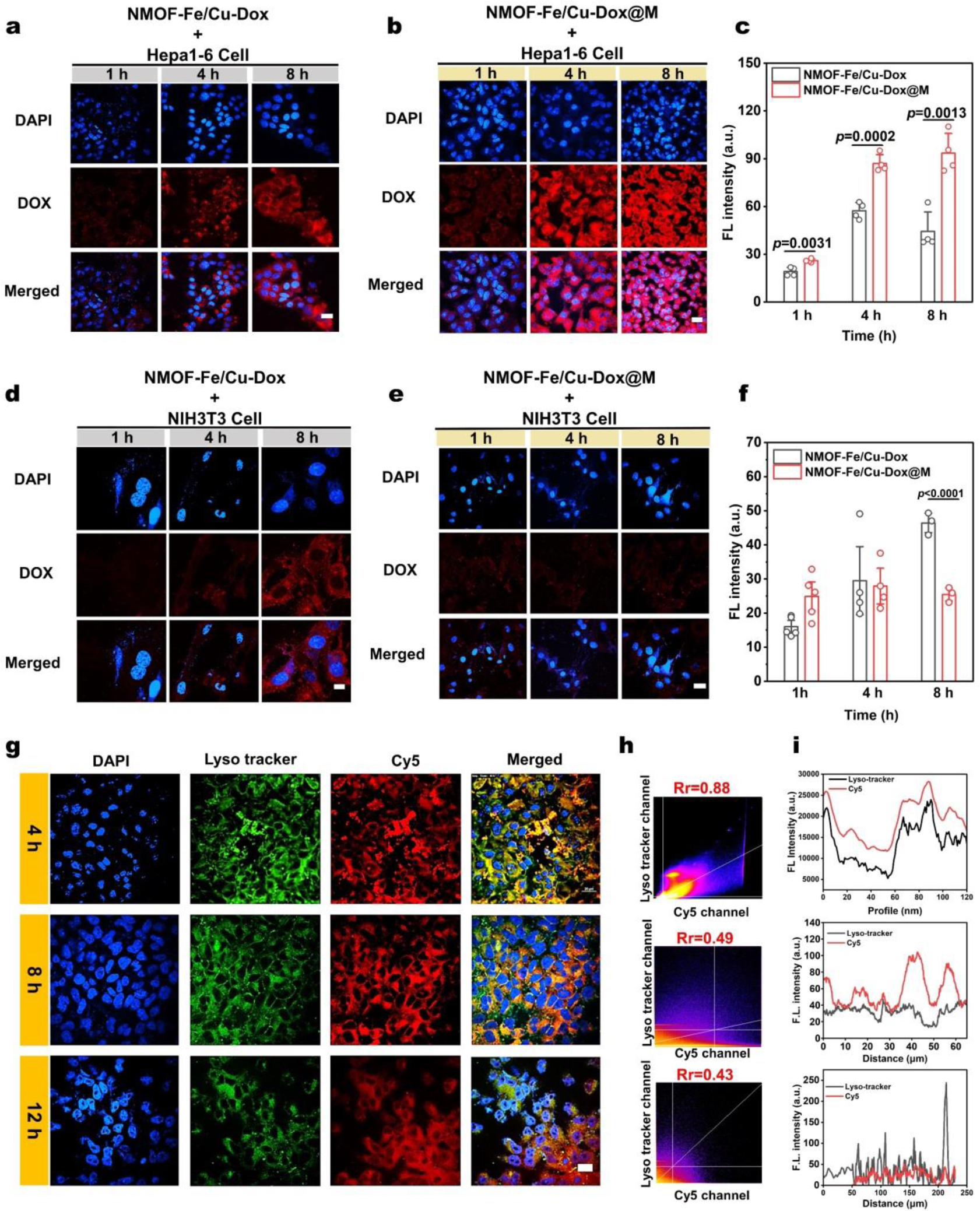
Evaluation of the homologous tumor-targeted delivery and subcellular localization of NMOF-Fe/Cu-Dox@M *in vitro*. **a**, **b**, Confocal images of the uptake of the NMOF-Fe/Cu-Dox and NMOF-Fe/Cu-Dox@M by Hepa1-6 cells at 1 h, 4 h and 8 h, Scale bar: 20 μm. **c**. Relative fluorescence intensities of Hepa1-6 cells incubated with NMOF-Fe/Cu-Dox and NMOF-Fe/Cu-Dox@M for 1 h, 4 h and 8 h, respectively (*n*=4). **d**, **e**, Confocal images of the uptake of the NMOF-Fe/Cu-Dox and NMOF-Fe/Cu-Dox@M by normal NIH3T3 cells at 1 h, 4 h and 8 h, Scale bar: 20 μm. **f**, Relative fluorescence intensities of NIH3T3 cells incubated with NMOF-Fe/Cu-Dox and NMOF-Fe/Cu-Dox@M for 1 h, 4 h and 8 h, respectively (*n*=4). **g**, Evaluation of lysosomal evasion of NMOF-Fe/Cu-Cy5@M, Scale bar: 20 μm. **h**, **i**, Pearson’s correlation coefficient and plot profile analysis of LysoTracker Green co-localization with NMOF-Fe/Cu-Cy5@M. The red lines representing NMOF-Fe/Cu-Cy5@M showed trajectories that were roughly similar to the black lines representing LysoTracker Green, indicating significant co-localization.

Next, to determine the distribution of the nanocomposites within the organelles of Hepa1-6 cells, the Cy5 loaded NMOF-Fe/Cu@M (NMOF-Fe/Cu-Cy5@M) was constructed instead of Dox in order to track the cellular uptake of the nanocomposites as a result of the high fluorescence quantum yield of the Cy5. After co-incubating NMOF-Fe/Cu-Cy5@M with cells for 4 h, the cells were stained using a commercial mitochondria probe MitoTracker Green, to assess the colocalization efficiency of NMOF-Fe/Cu-Cy5@M with the mitochondria. As shown in Supplementary Fig. S17, the results demonstrated a clear overlap in fluorescence between NMOF-Fe/Cu-Cy5@M and MitoTracker Green, indicating effective nanoparticle accumulation in the mitochondria. A high Pearson’s co-localization coefficient (PCC) value of 0.75 was obtained, indicating a strong correlation between Cy5 and MitoTracker Green. This finding demonstrated the efficacy of the nanocomposites in targeting mitochondria.

Besides, we also used the LysoTracker Green to investigate the relationship between the NMOF-Fe/Cu-Cy5@M and lysosomes. As shown in Fig. 3g, the NMOF-Fe/Cu-Cy5@M also exhibited a high co-localization with lysosomes at 4 h with a PCC value of 0.88. As the incubation time was prolonged to 8 h and 12 h, a notable portion of the red fluorescence from the NMOF-Fe/Cu-Cy5@M moved to the cytoplasm (PCC values of 0.49 and 0.43 at 8 h and 12 h, Fig. 3h and Fig. 3i), and exhibited no localization of the nanoparticles with lysosomes, indicating efficient evasion of the nanoparticles from lysosomal entrapment.

### Anti-tumor efficacy of NMOF-Fe/Cu-Dox@M *in vitro*

Initially, we evaluated the cytotoxicity of NMOF-Fe/Cu-Dox@M towards the normal NIH3T3 and HUVEC cells. As shown in Supplementary Fig. S18, the NMOF-Fe/Cu-Dox@M exhibited negligible impact on the proliferation of normal cells, indicating its high biocompatibility and safety for further *in vitro* and *in vivo* therapy. Next, the anti-tumor efficacy was investigated on Hepa1-6 cells *in vitro*. We also examined cytotoxic effect of single dose of NMOF-Cu, NMOF-Fe and NMOF-Fe/Cu SAzyme on Hepa1-6 cells, respectively. Compared to the NMOF-Fe and NMOF-Cu groups, NMOF-Fe/Cu SAzyme exhibited an enhanced cytotoxic effect on Hepa1-6 cells, indicating its synergistic effect in tumor therapy (Supplementary Fig. S19). Subsequently, we investigated and compared the anti-tumor efficacy of NMOF, NMOF-Fe/Cu, NMOF-Fe/Cu-Dox and NMOF-Fe/Cu-Dox@M to the Hepa1-6 cells. As shown in Supplementary Fig. S20, after co-administration with Dox, the cell viability of Hepa1-6 cells decreased to ∼ 45% at 4 h. Additionally, the inhibition of Hepa1-6 cell viability by NMOF-Fe/Cu-Dox@M was dose-dependent, with an inhibition rate of over 80% when the material concentration reached 80 μg mL^-1^, possibly due to the homologous internalization of Hepa1-6 cells (Fig. 4a). Furthermore, the therapeutic efficiency of NMOF-Fe/Cu-Dox@M on Heap1-6 cells showed slightly better performance after an incubation time of 24 h (Fig. 4b). Moreover, considering the acidity of solid tumors in the TME (pH 6.5-6.9), we particularly investigated the therapeutic effect of NMOF-Fe/Cu-Dox@M at pH 6.5. The results showed that, under pH 6.5, the proliferation inhibition effect of NMOF-Fe/Cu-Dox@M on Hepa1-6 cells was similar to that at pH 7.4 (Supplementary Fig. S21). These results indicated that the therapeutic effect was not affected by the extracellular pH microenvironment and NMOF-Fe/Cu-Dox@M could inhibit the tumor cell proliferation in such a condition.

**Fig 4.**
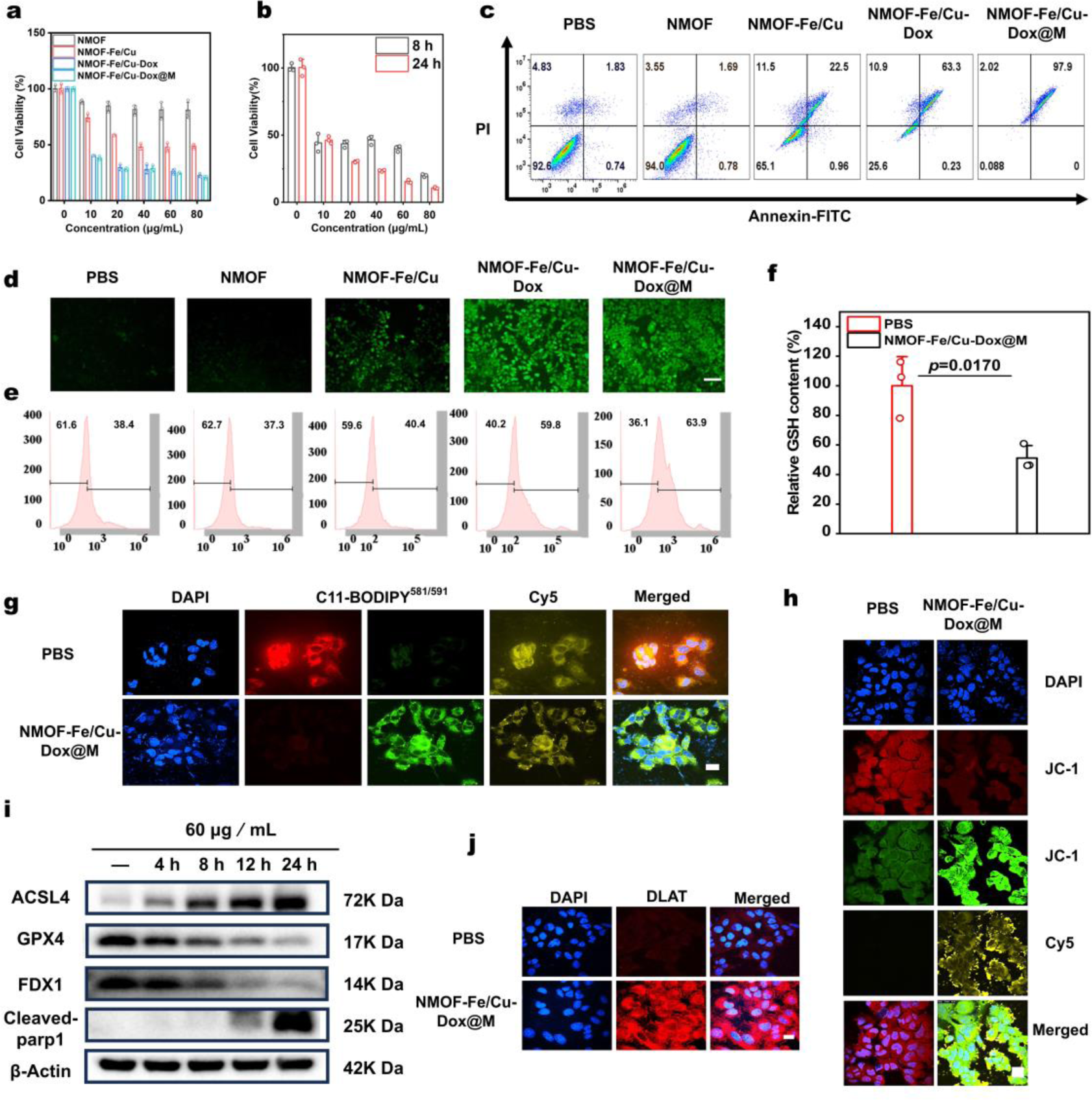
Evaluation of anti-tumor efficacy of NMOF-Fe/Cu-Dox@M *in vitro*. **a**, Cell viability of Hepa1-6 cells after incubated with NMOF, NMOF-Fe/Cu, NMOF-Fe/Cu-Dox and NMOF-Fe/Cu-Dox@M for 12 h (*n*=3). **b**, Cell viability of Hepa1-6 cells after incubated with NMOF-Fe/Cu and NMOF-Fe/Cu-Dox@M for 8 h and 24 h (*n*=3). **c**, Representative Annexin V-FITC/PI assay of Hepa1-6 cells after different treatments. **d**, Evaluation of ROS production on different treatments by DCFH-DA in Hepa1-6 cells, Scale bar: 20 μm. **e**, Corresponding flow cytometry analysis of ROS. **f**, Determination of the intracellular GSH levels in NMOF-Fe/Cu-Dox@M treated Hepa1-6 cells. Cells were treated for 12 h with the nanomedicine of 80 μg mL^-1^ and the GSH level was measured by a GSH/GSSG ratio kit (*n*=3). **g**. Lipid peroxide measured by C_11_-BODIPY^581/591^ staining in NMOF-Fe/Cu-Dox@M treated Hepa1-6 cells, Scale bar: 20 μm. **h**, Evaluation of the MMP by JC-1 in NMOF-Fe/Cu-Dox@M treated Hepa1-6 cells. Scale bar: 20 μm. **i**, Western blotting analysis of ACSL4, GPX4, FDX1 and Cleaved-parp1 expression in the NMOF-Fe/Cu-Dox@M treated Hepa1-6 cells at different time. **j**, DLAT fluorescence images of cancer cells after NMOF-Fe/Cu-Dox@M nanocomposites treated Hepa1-6 cells, Scale bar: 20 μm. Statistical significance was calculated by *t*-test.

Subsequently, the cell live/dead staining by Calcein-AM/PI was applied to evaluate the therapeutic efficiency of the NMOF-Fe/Cu-Dox@M towards the Hepa1-6 cells. In the PBS group and NMOF group, the Hepa1-6 cells showed brilliant green fluorescence (calcein-AM, green), indicating that NMOF did not produce cytotoxicity in cancer cells (Supplementary Fig. S22). In contrast, in the NMOF-Fe/Cu SAzyme group, a certain degree of dead Hepa1-6 cells was observed due to the chemodynamic therapy (CDT) induced by the POD-like and GPx-like enzyme activities. As expected, both of the NMOF-Fe/Cu-Dox and NMOF-Fe/Cu-Dox@M groups displayed excellent anti-tumor efficiency towards the Hepa1-6 cells. However, NMOF-Fe/Cu-Dox@M exhibited a little bit higher cytotoxicity than NMOF-Fe/Cu-Dox, possibly due to the cell membrane modification, which increased the uptake by the corresponding cancer cells. Additionally, we investigated the therapeutic effect of NMOF-Fe/Cu-Dox@M towards Hepa1-6 cells across different dosage levels. The results showed that the increasing of the NMOF-Fe/Cu-Dox@M had a positive cytotoxic effect on the viability of the Hepa1-6 cells (Supplementary Fig. S23). Furthermore, we examined the cell apoptosis by flow cytometry. As depicted in Fig. 4c, the experimental results were similar to those of the live/dead staining, with an increase in the proportion of apoptotic cells in the NMOF-Fe/Cu group, and a significant increase in apoptotic cells after incubation with NMOF-Fe/Cu-Dox@M, accounting for 69.8% of the total apoptotic cells, indicating that NMOF-Fe/Cu-Dox@M promotes apoptosis of cancer cells.

### Investigation of anti-tumor mechanisms

In previous reports, the ROS could not only directly trigger lipid LPO, but also modulate the genome in the condition of high oxidative stress to promote ferroptosis^46,51^. This implied that the production of large amounts of ROS could effectively drive ferroptosis. Despite ROS being capable of promoting ferroptosis, most ROS-targeted therapeutic approaches, such as photodynamic and chemodynamic therapies^41^, struggled to activate ferroptosis, due to excessive depletion of GSH alongside. Endogenous GSH, as an important substance maintaining the oxidative-reductive homeostasis of TME, could effectively eliminate the ROS at tumor sites and alleviate oxidative stress to some extent. Thus, elevating ROS levels at tumor sites while depleting endogenous GSH was expected to efficiently initiate tumor-specific ferroptosis. Since our NMOF-Fe/Cu SAzyme exhibited excellent inherent POD- and GPx-like enzyme activities, which could generate abundant ROS and simultaneously deplete GSH inside tumor cells, it was highly possible to induce the HCC cell death through the ferroptosis pathway. In order to prove these, we examined the involvement of ferroptosis and associated indicators, and explored the anti-tumor mechanism of NMOF-Fe/Cu-Dox@M against Hepa1-6 cells. By using dichloro-dihydro-fluorescein diacetate (DCFH-DA)^24^, an ROS probe, we initially examined the intracellular ROS in the HCC Hepa1-6 cells which were treated with a single dose of NMOF, NMOF-Fe/Cu, NMOF-Fe/Cu-Dox, NMOF-Fe/Cu-Dox@M and PBS, respectively. Fig. 4d showed that the NMOF-Fe/Cu, NMOF-Fe/Cu-Dox, and NMOF-Fe/Cu-Dox@M could effectively generate a high amount of intercellular ROS in the cancer cells, which could be due to excellent inherent POD-like enzyme activities of NMOF-Fe/Cu SAzyme. These findings were consistent with the results of the flow cytometry analysis (Fig. 4e). Moreover, escalating dosages of NMOF-Fe/Cu-Dox@M resulted in a progressive increase in intracellular ROS production in Hepa1-6 cells (Supplementary Fig. S24). Next, we delved deeper to explore whether the GPx-like activity of NMOF-Fe/Cu-Dox@M could deplete intracellular GSH levels. As described in Fig. 4f, 80 μg mL^-1^ of NMOF-Fe/Cu-Dox@M could deplete ∼50% intercellular GSH in the Hepa1-6 cells, suggesting the capability of NMOF-Fe/Cu-Dox@M to induce the cellular redox homeostasis. The simultaneous highly efficient generation of ROS and depletion of GSH could trigger the lipid LPO and induce the ferroptosis^52^. Then, we used the C_11_-BODIPY^581/591^ to evaluate the LPO in the Hepa1-6 cells^25^. The C_11_-BODIPY^581/591^ serves as an oxidation-responsive LPO probe, exhibiting a shift in maximum emission wavelength from 590 nm to 510 nm upon oxidation. As depicted in Fig. 4g, in NMOF-Fe/Cu-Dox@M group, the C_11_-BODIPY^581/591^ was oxidized to generate strong green fluorescence compared to the PBS group, indicating a highly LPO stimulated by NMOF-Fe/Cu-Dox@M in Hepa1-6 cells. However, the excessive ROS produced by NMOF-Fe/Cu-Dox@M might damage the mitochondrial membrane, resulting in ROS leakage and disruption of other organelles and biomacromolecules^53^. To assess the level of oxidative damage in HCC cells, we examined the mitochondrial depolarization by measuring the mitochondrial membrane potential (MMP) using a mitochondrial fluorescence probe, 5,5’,6,6’-tetrachloro-1,1’,3,3’-tetraethyl-imidacarbocyanine iodide (JC-1)^51^. The results indicate a loss of MMP in NMOF-Fe/Cu-Dox@M group compared to PBS group, leading to mitochondrial membrane depolarization and bright green fluorescence in the cytoplasm (Fig. 4h). These findings suggested that NMOF-Fe/Cu-Dox@M nanocomposites might induce ferroptosis in the HCC cells.

In general, overexpression of GSH in tumor cells could adapt to high levels of oxidative stress, consuming ROS and activating the GSH-related GPX4 expression to maintain cellular redox balance. The deactivation of GPX4 suggested that the defense and repair capability of the lipid antioxidants was largely inhibited, which would be beneficial for ferroptosis^54^. We then measured the GPX4 expression in the Hepa1-6 cells after the NMOF-Fe/Cu-Dox@M treatment. As expected, Western blotting results revealed both time- and concentration-dependent downregulation of GPX4 expression after NMOF-Fe/Cu-Dox@M treatment (Fig. 4i and Supplementary Fig. S25), which would favor the LPO generation. Meanwhile, we found that NMOF-Fe/Cu-Dox@M greatly boosted the expression levels of the Acyl-CoA synthetase long-chain family member 4 (ACSL4). Previous studies underscored the crucial role of ACSL4 in catalyzing the activation of arachidonic acid, through thioesterification with coenzyme A (CoA) to form arachidonoyl-CoA^52^. This arachidonoyl-CoA was subsequently incorporated into cellular membrane phospholipids and underwent oxidation catalyzed by lipoxygenase. This process of arachidonic acid released from phospholipase A2-catalyzed phospholipids, ultimately induced ferroptosis augmentation in HCC cells (Fig. 4i and Supplementary Fig. S25). These findings revealed the predominant impact of NMOF-Fe/Cu-Dox@M on various enzymes involved in regulating cellular redox processes, underscoring its potential to disturb redox homeostasis and trigger oxidative stress in HCC cells.

Afterwards, we investigated whether NMOF-Fe/Cu-Dox@M could induce cuproptosis, a novel mode of cell death initiated by copper ion binding to specific lipoylated molecules within the TCA cycle, a series of metabolic processes occurring within mitochondria^55^. During cuproptosis, excess Cu^+^ binds to lipoylated proteins in the TCA cycle, and results in remarkable aggregation of lipoyl synthase (LIAS)-mediated dihydrolipoamide dehydrogenase (DLAT), which indicated the extensive impact of disrupted metal homeostasis of cuproptosis on cellular processes. Herein, we examined the expression of the DLAT in HCC cells treated with NMOF-Fe/Cu-Dox@M. As depicted in Fig. 4j, the red fluorescence of DLAT in the NMOF-Fe/Cu-Dox@M group was highly strengthened compared to the PBS control group, suggesting that the NMOF-Fe/Cu-Dox@M successfully induced the DLAT aggregation to trigger cuproptosis. Moreover, the accumulation of Cu^+^ could disrupt the Fe-S cluster proteins and inactivate FDX1, potentially impacting downstream pathway of cuproptosis. Thus, we assessed the FDX1 expression levels in NMOF-Fe/Cu-Dox@M group by Western blotting. An effective down-regulation of FDX1 was observed (Fig. 4i and Supplementary Fig. S25), indicating the occurrence of cuproptosis. Finally, we evaluated the expression of the cell apoptosis marker, cleaved-parp1, which was found to be overexpressed in the NMOF-Fe/Cu-Dox@M group, revealing the efficient induction of the cancer cell death by ferroptosis/cuproptosis (Fig. 4i and Supplementary Fig. S25). In summary, our NMOF-Fe/Cu-Dox@M could synergistically initiate the intracellular ferroptosis/cuproptosis through the ROS storm, GSH/GPX4 depletion, ACSL4 activation and FDX1 inactivation, ultimately resulting in cell apoptosis.

### RNA-Seq analysis of the anti-tumor mechanism

To further elucidate the mechanism of NMOF-Fe/Cu-Dox@M against tumors, we performed transcriptome analysis on Hepa1-6 cells following exposure to both medicated (60 μg mL^-1^, 12 h) and non-medicated conditions. Utilizing the criteria of |log2 Fold Change| ≥ 1 and *p* value < 0.05, we identified that 4717 genes exhibited significant differential expression. The clustered heatmap visually illustrates an inconsistency in the gene expression disparities induced by the NMOF-Fe/Cu-Dox@M compared to the PBS group (Fig. 5a). Moreover, volcano plot of all differentially expressed genes revealed a down-regulation of the ferroptosis-associated gene GPX4 and an up-regulation of Steap3^56^. Concurrently, essential genes implicated in cuproptosis, i.e., FDX1, DLAT and DLST, demonstrated a decrease in expression. These observations are consistent with the above experimental outcomes (Fig. 5b). GO enrichment analyses showed a considerable enrichment of the pathways associated with the antitumor effects of NMOF-Fe/Cu-Dox@M, such as the regulation of developmental growth pathway and metal ion transmembrane transporter activity pathway (Fig. 5c). Ferroptosis and cuproptosis necessitate the accumulation of iron and copper ions within tumor cells, where they act at specific sites. Utilizing single-sample gene set enrichment analysis (ssGSEA) scoring, we identified pronounced differences in the expression of genes associated with the regulation, transport, binding, and homeostasis of these ions between NMOF-Fe/Cu-Dox@M and PBS groups. These findings suggest that NMOF-Fe/Cu-Dox@M disrupted the homeostasis of iron and copper ions in tumor cells, thereby facilitating the induction of ferroptosis and cuproptosis. Besides, cuproptosis, apart from its correlation with protein lipoylation in mitochondrial respiration, encompasses the loss of iron-sulfur cluster proteins in the respiratory chain. Our ssGSEA scoring revealed a discernible down-regulation in the expression of Fe-S cluster-related genes within tumor cells following drug administration (Fig. 5d).

**Fig 5.**
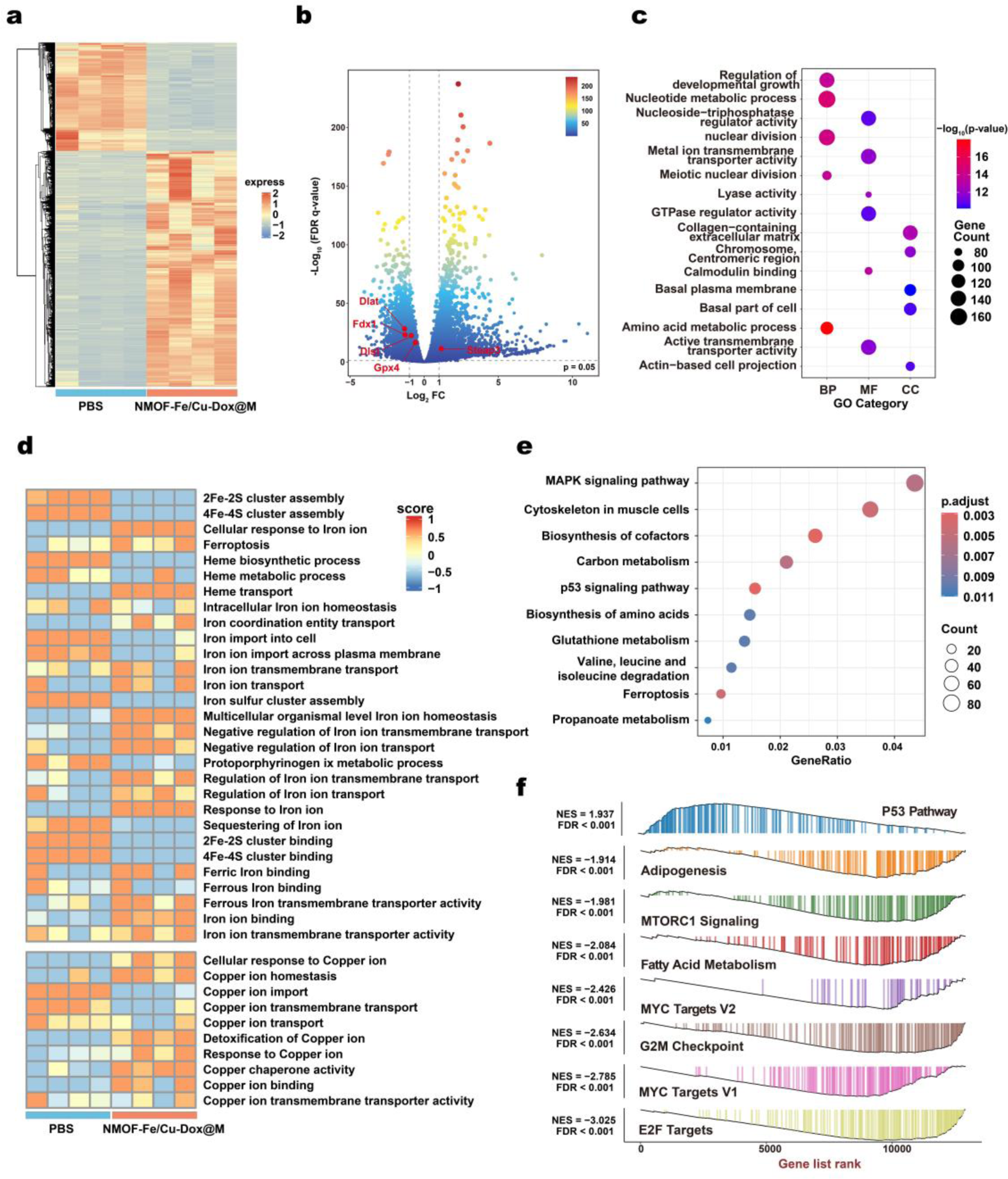
Investigation of the anti-tumor mechanism through the RNA-Seq analysis. **a**, The clustered heatmap of gene expression disparities induced by NMOF-Fe/Cu-Dox@M compared to the PBS groups. **b**, Volcano plot of differentially expressed genes. **c**, The GO enrichment analysis of the pathways associated with the antitumor effects by NMOF-Fe/Cu-Dox@M treatment. **d**, ssGSEA scoring of the iron and copper ion associated pathways. **e**, KEGG analysis of the ferroptosis pathway as well as the enrichment of both the p53 pathway and cell cycle-related pathways. **f**, GSEA of the p53 anti-cancer pathway and pertinent growth-proliferation-associated signaling pathways.

In the Kyoto Encyclopedia of Genes and Genomes (KEGG) analysis, we observed that, subsequent to NMOF-Fe/Cu-Dox@M treatment, activation of the ferroptosis pathway was accompanied by the enrichment of both the p53 pathway and cell cycle-related pathways (Fig. 5e). Following this, a Gene Set Enrichment Analysis (GSEA) was conducted, revealing activation of the p53 anti-cancer pathway after drug administration. Additionally, pertinent growth-proliferation-associated signaling pathways, such as MYC, mTORC1, G2M, and E2F, exhibited down-regulation consistent with *in vitro* cellular experiments (Fig. 5f). In previous studies, the inhibition of ferroptosis by both fat synthesis and fatty acid beta-oxidation had been established^17^. Through GSEA, we noted a down-regulation of pathways associated with adipogenesis and fatty acid metabolism. These observations also substantiated the capacity of NMOF-Fe/Cu-Dox@M to induce ferroptosis in tumor cells^55^. In summary, our meticulous analysis of the RNA-seq results reaffirms the capability of NMOF-Fe/Cu-Dox@M to elicit both ferroptosis and cuproptosis in tumor cells, concurrently suppressing pathways associated with growth and proliferation.

### *In vivo* anti-tumor efficiency of NMOF-Fe/Cu-Dox@M

Encouraged by the excellent therapeutic performance of NMOF-Fe/Cu-Dox@M against the HCC Hepa1-6 *in vitro*, we conducted the anti-tumor experiments *in vivo* (Fig. 6a). We first evaluated the biocompatibility and hemolysis of NMOF-Fe/Cu-Dox@M, and the results revealed that it possessed good biocompatibility and low hemolysis (< 5%, Supplementary Fig. S26). Next, we established the Hepa1-6-Luc subcutaneous HCC tumor xenografts in the C57BL/6 mice to evaluate the anti-tumor efficacy. Initially, we utilized fluorescence dye labeled NMOF-Fe/Cu-Dir@M, obtained by substituting Dox with Dir, to track the accumulation of the nanodrug at the tumor site and the distribution with *in vivo* fluorescence imaging. As shown in Fig. 6b, the fluorescence signals from NMOF-Fe/Cu-Dir@M were observed *in vivo* after 2 h injection, and reached its peak at 12 h, indicating the effective accumulation of NMOF-Fe/Cu-Dir@M in the tumor sites. In contrast, the fluorescence from the NMOF-Fe/Cu-Dir was remarkably weaker than that of NMOF-Fe/Cu-Dir@M, suggesting that the homologous tumor cell membrane was advantageous to enhance the uptake of the nanodrug by the tumor cells. Meanwhile, the NMOF-Fe/Cu-Dir@M still displayed stronger fluorescence after the post-injection of 24 h, indicating the cell membrane greatly prolonged retention of the nanodrug at the tumor site. These findings were consistent with the results of the previous cell experiments *in vitro* (Fig. 3a).

**Fig 6.**
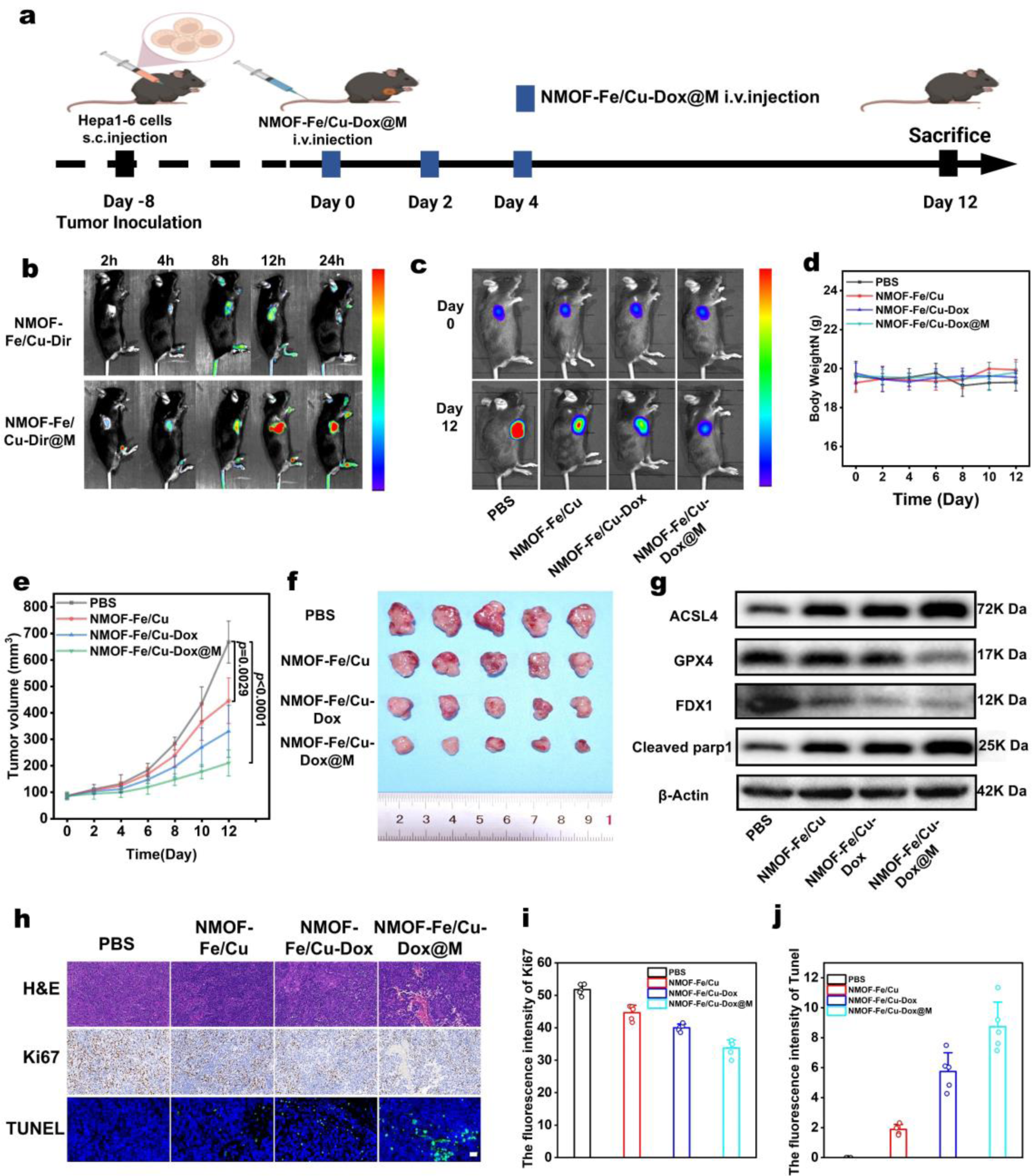
Evaluation of *in vivo* antitumor effect of NMOF-Fe/Cu-Dox@M on Hepa1-6-Luc subcutaneous mice tumor model. **a**, Schematic illustration of the experimental schedule *in vivo.* **b**, Biodistribution of NMOF-Fe/Cu-Dir and NMOF-Fe/Cu-Dir@M in tumors at different time points *in vivo*. **c**, The fluorescence images of tumor *in vivo* after different treatments on day 0 and day 12. **d**, Body-weight curves of mice (*n*=5). **e**, Tumor weight with different treated groups over time. **f**, Images of dissected tumors after different treatments at the end of the antitumor study (*n*=5). **g**, Western blotting analysis of the related proteins expression level in tumor tissues with different treatments. **h**, H&E, Ki67 and TUNEL staining of tumors with different treatment conditions, scale bar: 50 μm. **i, j,** The fluorescence intensity of Ki67 and TUNEL after various treatments (*n*=5).

Furthermore, we observed partial accumulation of the NMOF-Fe/Cu-Dir@M in the liver and lungs among main organs, likely attributed to clearance by the reticuloendothelial system (Supplementary Fig. S27).

Considering satisfactory safety, excellent targeting ability, and high uptake efficiency, we then evaluated the anti-tumor efficiency of NMOF-Fe/Cu-Dox@M *in vivo*. The NMOF-Fe/Cu-Dox@M and its controls, including the PBS, NMOF-Fe/Cu and NMOF-Fe/Cu-Dox, were subjected into the mice through the intravenous injection on days 0, 2, and 4, respectively (Fig. 6a). During the treatments, the NMOF-Fe/Cu-Dox@M group exhibited the strongest inhibition of tumor growth among all the studied groups (Fig. 6e), which was consistent with the fluorescence imaging results of the tumors in mice on day 12 *in vivo* (Fig. 6c, Supplementary Fig. S28). As depicted in Fig. 6d, almost no changes in body weights were observed throughout the treatment. The tumor growth inhibition index (TGI) of the NMOF-Fe/Cu-Dox@M group was calculated to be 60.54%. Meanwhile, we observed that the NMOF-Fe/Cu-Dox could also delay tumor growth but with a low TGI (28.33%) (Fig. 6e). At the end of the treatments, the mice were euthanized, and tumor tissues were dissected from the mice of all the groups for observation. The results indicated that the tumor tissues in the NMOF-Fe/Cu-Dox@M group were the smallest, further confirming the excellent therapeutic effect of NMOF-Fe/Cu-Dox@M (Fig. 6f).

We further extracted cells from tumor tissues in all groups and investigated the relevant protein expression via Western blotting analysis in order to check the ferroptosis and cuproptosis in mice (Fig. 6g). Compared to the other three groups, the ferroptosis-related phospholipid peroxidation, ACSL4, was obviously elevated, while the GSH-associated GPX4 protein was decreased in the NMOF-Fe/Cu-Dox@M group, indicating efficient activation of the ferroptosis pathway during treatment. Meanwhile, the upstream regulatory factor, FDX1 involved in the cuproptosis pathway, was markedly declined, suggesting that the NMOF-Fe/Cu-Dox@M also triggered cuproptosis pathway. The synergistic induction of ferroptosis and cuproptosis by NMOF-Fe/Cu-Dox@M enhanced the death of cancer cells, as evidenced by increased cleaved-parp1, a fragment of the full-length PARP1 protein resulting from caspase cleavage during apoptosis. Moreover, we utilized hematoxylin and eosin (H&E), Ki67 and terminal deoxynucleotidyl transferase dUTP nick end labeling (TUNEL) staining to analyze the tissues at the tumor sites. H&E staining assay could characterize the tumor tissue damage and the results demonstrate that the NMOF-Fe/Cu-Dox@M group exhibited the most tumor tissue damage, confirming the high therapeutic effect of the NMOF-Fe/Cu-Dox@M nanocomposites (Fig. 6h). Ki67, a widely utilized biomarker in cancer research for assessing cell proliferation and prognosis^57^, showed low expression in the NMOF-Fe/Cu-Dox@M group, indicating the efficacy of this nanodrug on inhibiting cancer cell proliferation and tumor growth (Fig. 6i). Correspondingly, the TUNEL staining demonstrated massive cancer cell apoptosis (TUNEL positive, green fluorescence) in the NMOF-Fe/Cu-Dox@M group compared to the other three groups, confirming the efficient induction of cancer cell apoptosis by NMOF-Fe/Cu-Dox@M (Fig. 6j). Finally, we stained the main organs of mice with H&E to assess the biocompatibility and safety of the different treatments. Analysis of heart, liver, spleen, lung, and kidney tissues revealed no apparent inflammation or organ abnormalities (Supplementary Fig. S29), suggesting minimal cytotoxicity of the NMOF-Fe/Cu-Dox@M.

Previous studies have reported that the peroxidized phospholipids produced during tumor ferroptosis might act as “eat me” signals to macrophages, promoting their polarization towards a pro-inflammatory state and thereby altering the tumor immune microenvironment^17^. However, the impact of ferroptosis on immune cells is still not well comprehended^27, 58^. Based on this, we utilized flow cytometry to examine the proportion and infiltration of intra-tumoral myeloid cells in PBS and NMOF-Fe/Cu-Dox@M groups. As shown in the Supplementary Fig. S30-S31, the proportion of M2 macrophages within the myeloid cell population decreased from 27.7% to 22%, suggesting that the treatment might alter the tumor immune microenvironment by modulating macrophage polarization.

### Evaluation of enhanced immune therapy of NMOF-Fe/Cu-Dox@M

Since the NMOF-Fe/Cu-Dox@M has been verified to efficiently induce the ferroptosis and cuproptosis, we proceeded to investigate whether it could modulate the TME to activate immune responses, thereby promoting the transformation of “cold” tumors into “hot” one. Initially, we investigated the macrophage re-polarization and anti-tumor immunity *in vitro*. In this case, we chose the conditioned media collected from blank Hepa1-6 cells and Hepa1-6 cells co-incubated with NMOF-Fe/Cu-Dox@M (60 μg mL^-1^, 12 h), termed as CM and CM_Hepa1-6_, respectively, to stimulate the macrophage cells and study their polarization. The RWA264.7 cells in the M0 state were cultured in both CM and CM_Hepa1-6_ for 48 h, and the subsequent polarization status was analyzed by flow cytometry (Fig. 7a and Supplementary Fig. S32). Under the culture condition of CM without drug treatment, 91.0% of RAW264.7 cells remained undifferentiated M0 status. However, we found only 17.9% of RAW264.7 cells remained M0 status under the CM_Hepa1-6_ treatment, which demonstrated that NMOF-Fe/Cu-Dox@M could remarkably activate the M0 macrophage polarization. Moreover, we observed an obvious increase in the proportion of M1 macrophages, which rose from 2.57% to 78.1%, along with a decrease in the proportion of M2 macrophages from 5.15% to 0.54%. These findings underscored the potent effect of NMOF-Fe/Cu-Dox@M in driving macrophages towards the M1 polarization, which could reverse the immunosuppression TME, stimulate the “cold” tumor to “hot” one and enhance the immune checkpoint therapy (Fig. 7b).

**Fig 7.**
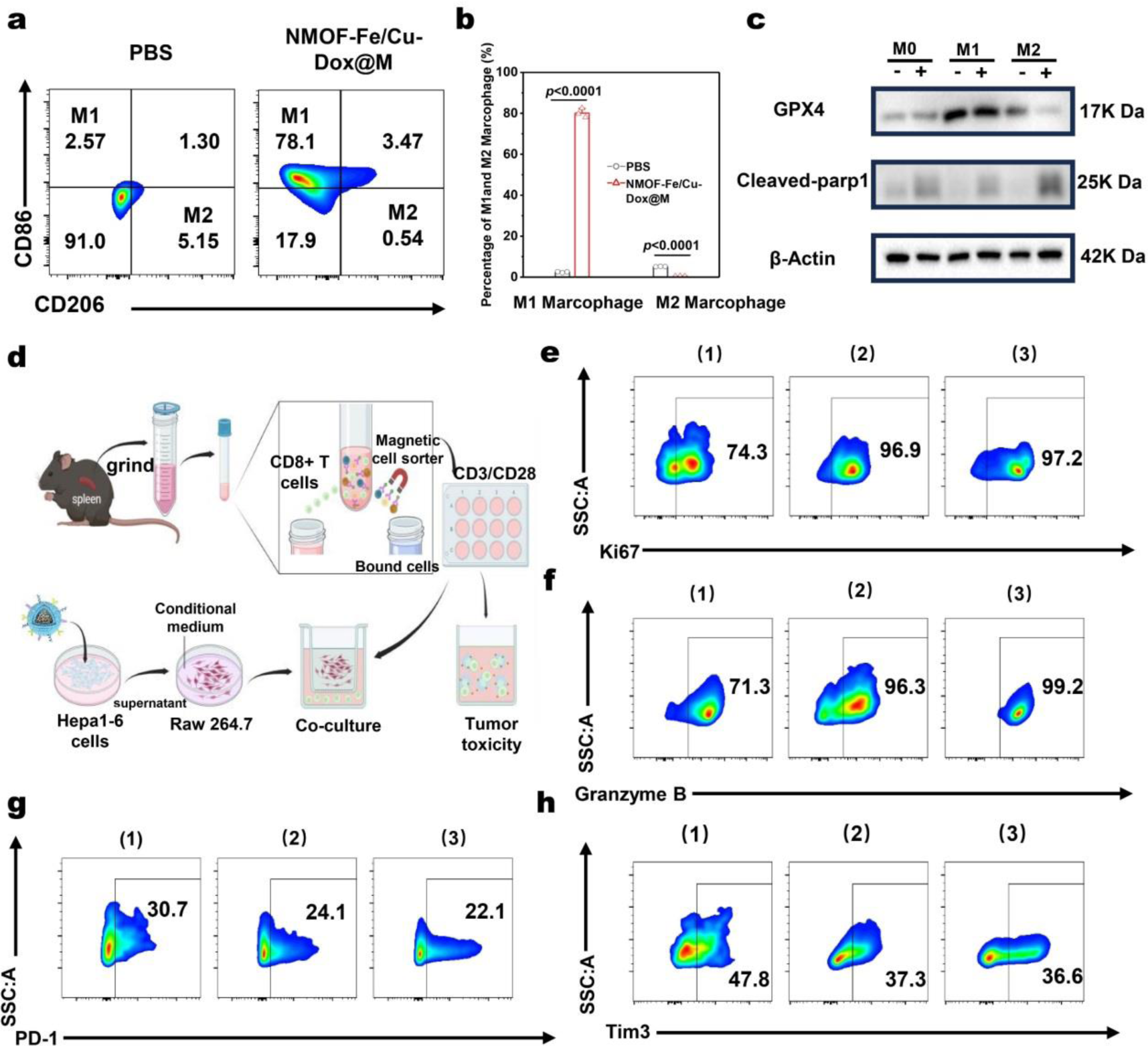
NMOF-Fe/Cu-Dox@M induced macrophage re-polarization via ferroptosis and anti-tumor immunity *in vitro*. **a**. Polarization status of M0 RAW264.7 cells after being cultured with CM and CM_Hepa1-6_. **b**, Corresponding percentages of M1 and M2 macrophages after the NMOF-Fe/Cu-Dox@M treatment (*n*=3). **c**. The ferroptosis and apoptotic phenotypes in RAW264.7 cells in M0, M1, and M2 states cultured with CM and CM_Hepa1-6_, examined by Western blotting. **d**, Schematic illustration of the impact of NMOF-Fe/Cu-Dox@M-induced ferroptosis on immune T cells *in vitro*. **e**, **f**, **g**, **h**, Representative flow cytometric plots of Ki67^+^, Granzyme B^+^, PD-1^+^ and Tim3^+^ on immune T cell surface after CM (1), CM_Hepa1-6_ (2) and CD3/CD28 (3) groups, respectively. Statistical significance was calculated by *t*-test.

We then examined the impact of ferroptosis induced by NMOF-Fe/Cu-Dox@M on macrophages. Some previous investigations have demonstrated that, compared to M2 macrophages, M1 macrophages overexpress inducible nitric oxide synthase (INOS) and exhibit higher tolerance to ferroptosis^59^. Thus, we employed the lipopolysaccharide (LPS) and IFN-γ to stimulate the RAW264.7 cells to obtain the M1 macrophages, as well as interleukin-4 (IL-4) and interleukin-10 (IL-10) to obtain M2 macrophages. Subsequently, M0, M1, and M2 macrophages were treated with or without NMOF-Fe/Cu-Dox@M (60 μg mL^-1^, 12 h) and the relevant indicators of ferroptosis were examined. As Fig. 7c depicted, we found no obvious change in GPX4 expression in M0 and M1 macrophages, indicating the minimal influence on these macrophages by ferroptosis. However, the M2 macrophages exhibited a notable decrease in GPX4 levels, revealing that NMOF-Fe/Cu-Dox@M could efficiently induce the ferroptosis in M2 macrophages. As expected, a noticeable increase in apoptosis marker (cleaved-parp1) was detected on NMOF-Fe/Cu-Dox@M treated M2 macrophages, which further proved that NMOF-Fe/Cu-Dox@M could efficiently induce the M2-phenotype macrophages apoptosis. These findings underscored that NMOF-Fe/Cu-Dox@M could induce the pro-tumor M2 macrophages death via the ferroptosis, increase the frequency ratio of the anti-tumor M1 macrophages, reverse the immunosuppressed TME and enhance the cancer treatment efficiency.

Apart from above experiments, we investigated the impact of ferroptosis on immune cells and its role in modulating the immune cells *in vitro*. This, in turn, caused the activation and enhancement of T cell-mediated anti-tumor immunotherapy. As Fig. 7d depicted, the immune cells were extracted from the spleens of normal C57 BL/6 mice and CD8^+^ T cells were selected by magnetic isolation. These CD8^+^ T cells were then co-cultured with RWA264.7 cells treated with CM and CM_Hepa1-6_ for 48 h, followed by flow cytometric analysis. It was worth noting that the CD8^+^ T cells were continuously activated with CD3 and CD28 antibodies, which were then employed as the positive group to assess the level of activation and proliferation^60^. The flow gating strategy for analysis of the CD8^+^ T cells in tumors was depicted in the Supplementary Fig. S32. The elevated cytotoxicity of Granzyme B^+^ levels in CD8^+^ T cells, along with the proliferation activity indicated by the Ki67^+^ biomarker during treatment, suggested the activation of more CD8^+^ T cells by immunotherapy, consequently further enhancing their ability to attack tumors^61^. Thus, we evaluated the Ki67^+^ and Granzyme B^+^ levels in these CD8^+^ T cells. Compared to the CM group (1), the CM_Hepa1-6_ group (2) and CD3/CD28 group (3) exhibited a higher proportion of Ki67^+^/CD8^+^ T cells and Granzyme B^+^/CD8^+^ (Fig. 7e and 7f), indicating the elevated proliferation activity and cytotoxicity of CD8^+^ T cells. These findings demonstrated the increased proliferation activity of CD8^+^ T cells and elevated cytotoxicity in CD8^+^ T cells during the NMOF-Fe/Cu-Dox@M treatment, suggesting that more CD8^+^ T cells were activated by immunotherapy and further promoted to attack tumors.

In addition, we checked the expression of PD-1^+^ and Tim3^+^ in CD8^+^ T cells, which were two immune checkpoint molecules and played crucial roles in regulating T cell function. Overexpression of these molecules might lead to functional exhaustion of T cells, resulting in a loss of their ability to effectively attack tumor cells. On the contrary, the reduction of PD-1^+^ and Tim3^+^ expression in CD8^+^ T cells could be interpreted as the restoration of T cell function or alleviation of their exhaustion, thereby enhancing the immune response against cancers. We then measured the proportion of PD-1^+^/CD8^+^ and Tim3^+^/CD8^+^ T cells, and observed a decrease in these biomarker-expressing T cells (Fig. 7g and 7h), implying a reduction in T cell exhaustion. Interestingly, we noticed that the activation and regulation level of T cells in the CM_Hepa1-6_ group was comparable to that of CD3/CD28 group, demonstrating the highly efficient immune activation of NMOF-Fe/Cu-Dox@M (Supplementary Fig. S33). Subsequently, we assessed the cell viability under the T cells activation and found that the T cells in the CM_Hepa1-6_ group exhibited significantly enhanced immunotherapy (Supplementary Fig. S34).

In summary, our NMOF-Fe/Cu-Dox@M could not only induce ferroptosis/cuproptosis in Hepa1-6 cells but also modulate macrophage polarization to M1 macrophages and exhaust M2 macrophages, promoting immune activation and stimulating immune T cells for synergistic immunotherapy. Expectedly, NMOF-Fe/Cu-Dox@M would be promising for highly efficient eradication of tumor cells.

### Combination ferroptosis/cuproptosis with immune checkpoint therapy impeding the progression of HCC

Based on above results, we found that the treatment reversed macrophage polarization towards a pro-tumor state both *in vitro* and *in vivo*. Notably, *in vitro* experiments revealed that the antitumor function of CD8^+^ T cells was boosted as macrophages underwent repolarization. Herein, we were conducting further investigations to determine its potential to reconfigure TME *in vivo* and enhance the sensitivity of immune checkpoint therapy, using the Hepa1-6-Luc subcutaneous HCC mouse model continuously. As shown in the experimental schedule (Fig. 8a), we added the αPD-1 monotherapy group (100 μg, intraperitoneal injection) and a combination group (NMOF-Fe/Cu-Dox@M and αPD-1) into the study. As depicted in Fig. 8b, 8c and Supplementary Fig. S35, compared to the PBS group, the αPD-1 group exhibited considerable anticancer activity, consistent with findings from other studies^62^. Importantly, NMOF-Fe/Cu-Dox@M demonstrated comparable efficacy to αPD-1 treatment, while the combination therapy group showed a markedly superior anti-tumor effect to either of the above two monotherapies. From the perspective of tumor progression, this trend is conspicuously apparent (Fig. 8d). Moreover, the H&E staining and immunofluorescence staining of subcutaneous tumors further validated that the combination therapy was more effective in inhibiting tumor proliferation and inducing tumor apoptosis (Supplementary Fig. S36 and S37). Furthermore, there was no notable *in vivo* toxicity, as evidenced by the lack of obvious changes in mouse body weight, blood biochemical markers and organ H&E staining (Supplementary Fig. S38 and S39). As depicted in Supplementary Fig. S40, the NMOF-Fe/Cu-Dox@M and αPD-1 did not induce the alterations in the indices of liver function, renal function indicator and blood cell index in comparison to the PBS control group.

**Fig 8.**
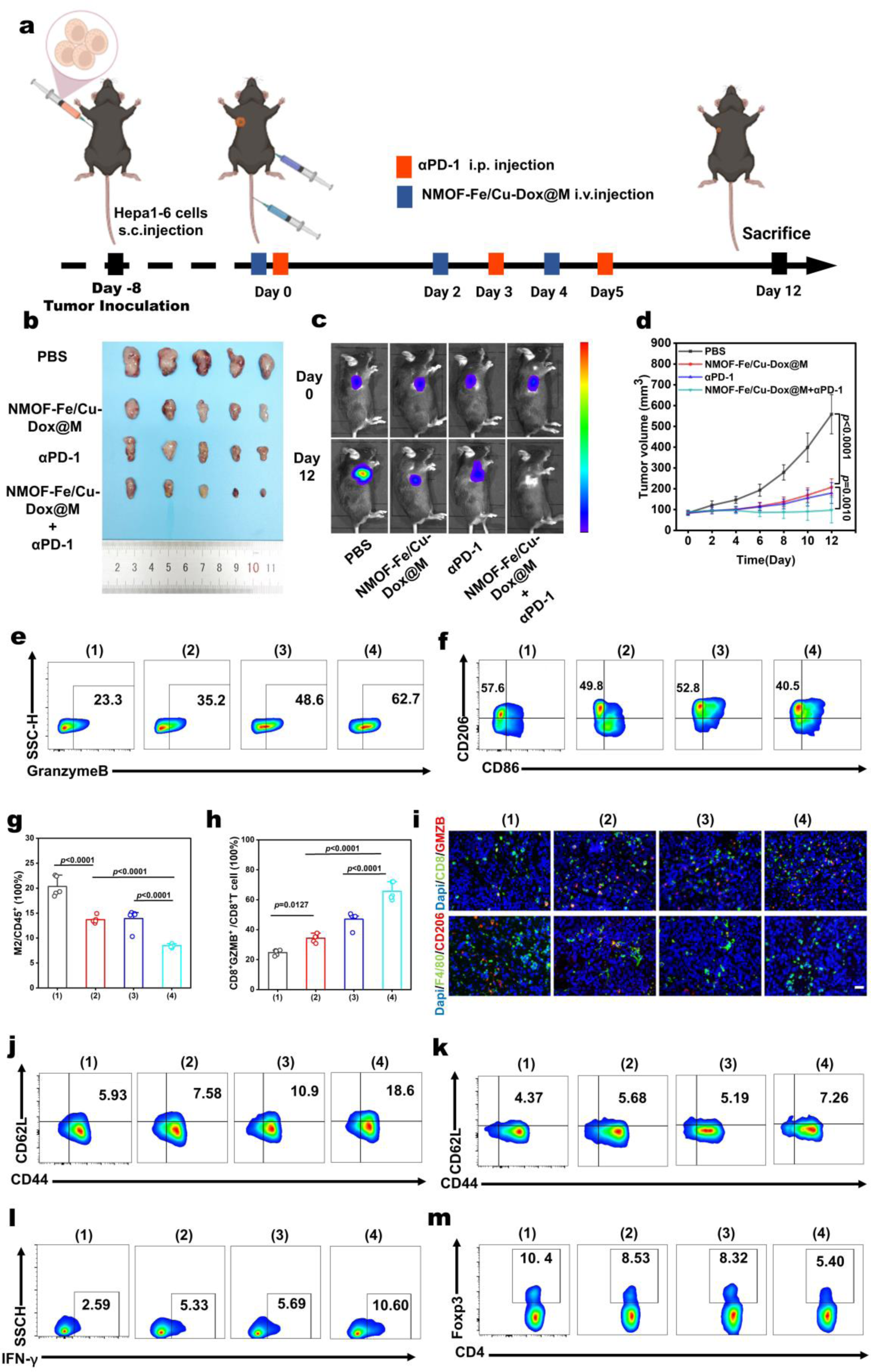
*In vivo* antitumor effect of NMOF-Fe/Cu-Dox@M combined with αPD-1 therapy. **a**, Schematic illustration of the antitumor and immunotherapy experimental schedule *in vivo*. **b**, Photos of tumors at the end of *in vivo* antitumor and immunotherapy study (*n*=5). **c**, Bioluminescence images of tumors after antitumor and immunotherapy. **d**, **e**, Tumor volume and body weight variation in different treatments. **e**, **f**, **g**, **h**, M2 macrophage and Granzyme B^+^ T cell in Hepa1-6 subcutaneous mice tumor model after different treatments (*n*=5). **i**, Tumor tissue immunofluorescent staining, scale bar: 50 μm. **j**, Representative flow cytometric plots of CD8^+^ T cells in tumors after different treatments. **k**, Representative flow cytometric plots of Treg cells and in tumors of mice after different treatments. **l**, Representative flow cytometric plots of CD4^+^/IFNγ^+^ T cells in tumors after different treatments. **m**, Representative flow cytometric plots of CD4^+^ Tcm cells in tumors after different treatments. (1) PBS; (2) NMOF-Fe/Cu-Dox@M; (3) αPD-1; (4) NMOF-Fe/Cu-Dox@M+αPD-1. Statistical significance was calculated by *t*-test.

Next, we conducted flow cytometric analysis to investigate potential alterations in the immune TME that might underlie the sensitization of immunotherapy by NMOF-Fe/Cu-Dox@M. We extracted immune cells from tumor tissues and performed flow cytometric analysis (Supplementary Fig. S41). The flow cytometric results indicated that there were no obvious changes in the quantity of infiltrating CD8^+^ and CD4^+^ T cells within the tumor. However, remarkable alterations were observed in the differentiation and effector markers of M2 macrophages cells (CD86^-^/CD206^+^). This phenomenon was consistent with our expectations, as Hepa1-6-Luc subcutaneous HCC mouse model had the “hot” TME, wherein a large number of macrophages had already differentiated towards M1 upon initial treatment^63^. Compared to the individual therapy groups of (1) PBS, (2) NMOF-Fe/Cu-Dox@M and (3) αPD-1, the combination therapy group of (4) NMOF-Fe/Cu-Dox@M+αPD-1 exhibited the lowest proportion of M2 macrophages (CD45^+^, 8.9%), which was prominently lower than that in the PBS (22.6%), NMOF-Fe/Cu-Dox@M (13.7%) and αPD-1 (14.9%) groups (Fig. 8e and 8f). Notably, compared to the individual monotherapy groups, the combination therapy group exhibited a significant increase in granzyme B^+^/CD8^+^ (Fig. 8g and 8h), indicating the ability of NMOF-Fe/Cu-Dox@M+αPD-1 to induce the infiltration of memory T cells and reverse the TME for the enhanced immune checkpoint therapy. In addition, the immunofluorescence staining of mouse tumor tissues also revealed a significant increase in the proportion of granzyme B^+^/CD8^+^ T cells in the group of NMOF-Fe/Cu-Dox@M+αPD-1, along with a noticeable decrease in M2 macrophages (F4/80^+^/CD206^+^) in the TME, consistent with the results of *in vivo* and *in vitro* experiments (Fig. 8i).

Moreover, the proportion of CD44^+^/CD62L^+^ T cells within both CD8^+^ and CD4^+^ T cells populations remarkably increased after NMOF-Fe/Cu-Dox@M or αPD-1 treatments, as well as the combination therapy of both. This indicated the ability of NMOF-Fe/Cu-Dox@M to induce infiltration capability of the memory T cells, specially CD8^+^ central memory T cells (CD8^+^ Tcm, Fig. 8j and Supplementary Fig. S42a) and CD4^+^ T helper memory cells (CD4^+^ Thm, Fig.8k and Supplementary Fig. S42b), and to reprogram the TME to enhance the ICB therapy. Furthermore, there was a notable increase in IFN-γ^+^/CD4^+^ T cells, indicating that NMOF-Fe/Cu-Dox@M could enhance antitumor efficiency by modulating T cell differentiation (Fig. 8l and Supplementary Fig. S42c). Meanwhile, the proportion of classical pro-tumor Treg cells (CD4^+^/Foxp3^+^) decreased after NMOF-Fe/Cu-Dox@M+PD-1 combination treatment (Fig. 8m and Supplementary Fig. S42d). From the above *in vivo* experiments, we can see that NMOF-Fe/Cu-Dox@M could reduce the polarization of macrophages towards M2 and reprogram the TME to enhance antitumor efficiency *in vivo*. In summary, the NMOF-Fe/Cu-Dox@M exhibited a synergistic antitumor effect and enhanced the sensitivity of ICB therapy in HCC.

## Discussion

Cancer immunotherapy employs strategies to activate and harness the immune system to eradicate cancer. Among them, ICB exhibited exceptional anti-cancer responses in patients by targeting the cytotoxic T-lymphocyte-associated protein 4, PD-1, and PD-L1 molecules on immune T cells and cancer cells^1, 3, 6^. The effectiveness of ICB therapies is typically considered to rely on the presence of pre-existing anti-tumor T-cell immunity^8, 64^. However, in an immunologically “cold” tumor, such as HCC, the TME typically exhibits immunosuppressive characteristics, including high expression of tumor-associated myeloid cells (TAMCs), like myeloid-derived suppressor cells (MDSCs), tumor-associated macrophages (TAMs, M2 macrophages), regulatory T cells (Tregs), and immune checkpoint molecules (e.g., IL-6, TGF-β), which affect the efficacy of ICB therapy^2, 3, 59^. Meanwhile, inefficient T cell infiltration further hinders the effectiveness of ICB in cancer therapy^4, 6, 8^. To overcome these challenges, effective strategies, e.g., triggering immunogenic cell death, stimulating the immune system to boost the anti-tumor T cell response, and reversing the immunosuppressive TME, can be used to facilitate combination immunotherapy with ICBs to elicit a more robust anti-tumor immunity.

Recent evidence has shown that triggering redox dyshomeostasis, such as the generation of excessive ROS *in vivo*, could be potent to enhance the immune response^17, 29^. This discovery inspirates the development of deployable immunogenic treatment modalities as immunostimulatory adjuvants to enhance ICB therapy. Ferroptosis, a novel form of non-apoptotic cell death that operates via redox dyshomeostasis and lipid LPO, shows great potential in cancer treatment by inducing immunogenic tumor cell death and enhancing the immune response. However, the therapeutic efficacy of ferroptosis is generally hampered by tumor-intrinsic protective mechanisms associated with endogenous GSH or metabolites derived from the amino acid cysteine^65^. Thus, we thought to design nanomedicines that can induce ferroptosis and synergistically integrate with cuproptosis to modulate the TME by depleting the intrinsic antioxidants, and activating robust local immune responses to enhance ICB therapy.

Herein, we have developed a Fe^3+^/Cu^2+^ bimetallic ions functionalized UIO-67 NMOF SAzyme and loaded it with Dox, termed as NMOF-Fe/Cu-Dox, to achieve synergistic enzyme-catalytic therapy, chemotherapy and immunotherapy for HCC. As anticipated, the engineered NMOF-Fe/Cu-Dox@M showed remarkable efficacy in TME-responsive catalytic therapy by increasing cellular oxidative stress. This was achieved through a dual mechanism: enhanced ROS production with POD-like enzymatic activity and depletion of GSH levels with GPx-like enzymatic activity in HCC. The accumulated ROS induced LPO, triggering ferroptosis in the cancer cells. Simultaneously, the excessive Cu^2+^ in the NMOF-Fe/Cu-Dox@M can cause the aggregation of lipoylated protein DLAT and depletion of Fe-S cluster proteins, leading to cellular cuproptosis. Concurrently, the redox dyshomeostasis promoted LPO and GPX4 inactivation, enhancing the ferroptosis effect. The increased oxidative stress drove synergistic ferroptosis and cuproptosis, collectively causing mitochondrial dysfunction and contributing to a significant therapeutic impact. In summary, the redox dyschondrosteosis and excessive Fe³⁺/Cu²⁺ induced strong ferroptosis, which worked synergistically with cuproptosis and Dox-mediated apoptosis to effectively inhibit tumor growth.

In our design, the NMOF-Fe/Cu-Dox is camouflaged with a cancer cell membrane, which facilitates homologous targeting of cancer cells, enhances tumor penetration, and achieves immune evasion. The bioinspired approach utilizes natural cancer cell membrane coating technology, which efficiently delivers nanocomposites to desired sites due to natural intercellular interactions, thereby increasing therapeutic efficacy and safety^42, 66, 67^. Thus, our NMOF-Fe/Cu-Dox@M can cross biological barriers, avoid immune elimination, specifically target and accumulate in tumors *in vivo*, and exhibit exceptional safety characteristics. It is worth noting that this is important for ferroptosis-associated cancer therapy, as previous reports have demonstrated that ferroptosis can influence immune cells, such as myeloid-derived suppressor cells (PMN-MDSCs) in TME, which spontaneously die by ferroptosis and promote tumor growth^58^. This reminds us to be very cautious when using ferroptosis and cuproptosis-associated therapy to avoid affecting T cells, such as neoantigen-specific cytotoxic Tr1 CD4 T cells^68^. Fortunately, the cancer cell membrane camouflaging the nanocomposites allows us to circumvent this issue and ensures the safety of our approach.

Simultaneously, we found that the NMOF-Fe/Cu-Dox@M nanocomposites could induce the death of immunosuppressive M2 macrophages without affecting immunostimulatory M1 macrophages, thereby significantly increasing the M1/M2 macrophage ratio and enhancing the immune response. This process might work synergistically with mature dendritic cells (DCs) to enhance antigen processing and presentation to naïve T cells. As a result, NMOF-Fe/Cu-Dox@M treatment promoted robust tumor-infiltration by CD8^+^ T cells and increased their proliferation and functionality *in vivo*, as evidenced by the increased expression of Ki67^+^ and granzyme B^+^ on them. These CD8^+^ T cells could recognize and kill tumor cells, releasing additional tumor antigens and enhancing the immune response with the support of ICB therapy^8^. Conversely, the decrease in PD-1^+^ and Tim3^+^ expression on CD8^+^ T cells suggests a restoration of T cell function and a reduction in their exhaustion, thereby strengthening the immune response against cancers. Further combining NMOF-Fe/Cu-Dox@M treatment with the αPD-1 antibody effectively treated HCC, significantly enhanced therapeutic efficacy, activated local immune responses, and bolstered the impact of ICB therapy in a more pronounced way. This study not only presents novel SAzyme-based nanocomposites with remarkable enzymatic activities that can reverse the immunosuppressive TME and activate immunity for the cancer therapy, but also paves the way for broadening the biological applications of synergistic ferroptosis/cuproptosis with enhanced ICB for other cancers.

### Statistical information

All results are expressed as mean ± SD (standard deviation), and differences between groups were calculated through Student’s ***t***-tests. Differences with *P* values less than 0.05 were considered significant. The statistical details of the experiments are included in the figure captions. The Origin 2024b was used for statistical analysis.

## Data Availability

Source data are provided with this paper. The authors declare that all other data supporting the findings of this study are available within the paper, Supplementary Information or Source Data file. Source data are provided with this paper.

## Ethics declarations

Animal handling, surveillance, and experimentation were performed in accordance with and approval from the Animal Use Committee of Tongji Hospital of Huazhong University of Science and Technology (TJH-202310024).

## Acknowledgements

This work is supported by the National Natural Science Foundation of China (Grant Nos. 22174049, 82373834, 82172976, 82303628), the Program for HUST Academic Frontier Youth Team (Grant No. 2019QYTD09) and Natural Science Foundation of Hubei Province (Grant Nos. 2024AFA076). The authors also thank the Analytical and Testing Center of HUST, and Medical Sub Center of Analytical and Testing Center of HUST for the material characterization.

## Author Contributions

L.Y., M.P, H.H., F.L. Z.Z., X.Y. and L.X. conceived and designed the experiments. L.Y., M.P., H.H and F.L. conducted all experiments. L.Y., M.P., J.M. and C.X. assisted with protein assay, cell culture and *in vitro* assays. H.H., F.L. and S.N. helped with all *in vivo* studies. L.Y. and M.P. supported the biodistribution and safety studies. H.H. and F.L. assisted with tumor efficacy studies. L.Y., M.P., H.H., F.L. Z.Z., X.Y. and L.X. analyzed and discussed the data. L.Y. and H.H. interpreted the data and wrote the paper. Z.Z., X.Y. and L.X. wrote, reviewed and edited the paper.

## Supplementary Information

The online version contains supplementary material available at https://XXXXX

