## Supplementary for "Bioinspired Bimetallic Ions Functionalized MOF SAzyme Nanocomposites for Synergistic Ferroptosis/Cuproptosis-Enhanced Immune Checkpoint Therapy"

Contributions

#These authors contribute equally to this study.

Corresponding author

Correspondence to: Zhan-Guo Zhang[2]*, Xu Yu[1]*, Li Xu[1]*

.

### Supplementary methods

#### Materials

Ethanol, methanol, tetrahydrofuran (THF), N, N-dimethylformamide (DMF), $ZrCl_4$, dimethyl sulfoxide (DMSO), MB and TA were obtained from Sinopharm Chemical Reagent Co., Ltd (Shanghai, China). Doxorubicin hydrochloride (Dox), 2,2′-bipyridine-5,5′-dicarboxylic acid ($H_2BPY$), 2',7'-Dichlorodihydrofluorescein diacetate (DCFH-DA) and 3,3′,5,5′-tetramethylbenzidine (TMB) were acquired from Aladdin Chemical Reagent Co., Ltd. (Shanghai, China). $FeCl_3 \cdot 6H_2O$ and $Cu(NO_3)_2 \cdot 2H_2O$ were acquired from Aladdin Chemical Reagent Co., Ltd. (Shanghai, China). Membrane and Cytosol Protein Extraction Kit, phenylmethylsulfonyl fluoride (PMSF), RIPA solution, Calcein/PI Cell Viability/Cytotoxicity Assay Kit and Lyso-Tracker Green were purchased from Beyotime Biotechnology. FITC Annexin V Apoptosis Detection Kit I was purchased from BD Biosciences. JC-1 and Mito- Tracker Green were purchased from MedChem Express Co., Ltd. $C_{11}$-BODIPY$^{581/591}$ was purchased from Thermo Scientific Co., Ltd. DMEM/HIGH Glucose culture was purchase from Monad Biotech Co., Ltd. Immobilon-P PVDF membrane was purchased from Merck Millipore Ltd.

#### Characterizations

The content of Fe and Cu incorporated in the NMOF-Fe/Cu was determined by the inductively coupled plasma mass spectrometry (ICP-MS, PlasmaQuant® PQ 9000 Elite). The morphology of NMOF, NMOF-Fe/Cu, NMOF-Fe/Cu-Dox, NMOF-Fe/Cu-Dox@M were observed by a Field emission scanning electron microscope (SEM, MIRA 3, TESCAN Brno, s.r.o., Russia). Transmission electron microscopic (TEM) images were observed by a Talos F200X Field emission 603 microscope (FEI, Netherlands). Spherical Aberration Corrected Transmission Electron Microscope

(ACTEM) images were observed by a Thermo Fisher Titan Themis G2 60-300 microscope. Zeta potential analyzer (Brookhaven, USA) was utilized to determine the zeta potential. Thermal gravimetric analysis (TGA) of NMOF, NMOF-Fe/Cu and NMOF-Fe/Cu-Dox were performed by a TG 209 F1 Libra thermal analyzer (Netzsch, Germany). The state of single Fe and Cu and their complexes in NMOF-Fe/Cu, were verified by the X-ray photoelectron spectroscopy (XPS, Thermo Fischer, American).

**Experimental sections**

Synthesis of $Zr^{4+}$-NMOFs (NMOF): NMOF was prepared according to the previously reported literature with slight modification[1, 2]. In a typical procedure, 48.8 mg of $ZrCl_4$ (0.2 mM) and 46.6 mg of $H_2BPY$ (0.2 mM) were dissolved in 20 mL DMF, respectively. After treatment with ultra-sonification for 5 min, the two solutions were mixed into a 100 mL glass bottle, and then 930 μL of acetic acid was added into the mixture solution. The mixture solution was heated at 120 °C for 18 h. After cooling down to room temperature, the white solids were collected by centrifugation at 9500 rpm. Next, the obtained NMOF was washed with DMF, THF, ethanol, and deionized water, respectively. The purified NMOFs were dispersed in ethanol.

**Synthesis of NMOF-Fe/Cu:** For the synthesis of the NMOF-Fe/Cu, 10 mg of NMOF were added to 670 μL $FeCl_3 \cdot 6H_2O$ ethanol solution (1 M) and 330 uL $Cu(NO_3)_2 \cdot 2H_2O$ aqueous solution (1 M). The mixture solution was incubated on a rotary shaker for 4 h at room temperature. After washing with ethanol and deionized water for several times, the excess $FeCl_3 \cdot 6H_2O$ and $Cu(NO_3)_2 \cdot 2H_2O$ were removed by centrifugation and the NMOF-Fe/Cu was obtained.

**Synthesis of NMOF-Fe/Cu-Dox:** For Dox loading, 10 mg NMOF-Fe/Cu was dispersed in the Dox aqueous solution (10 mg·mL$^{-1}$, 1 mL) with stirring for 12 h. After that, NMOF-Fe/Cu-Dox were collected by centrifugation (10,000 rpm, 5 min) and further dispersed in ultrapure water.

**Preparation of Hepa1-6 Cell Membrane:** The cell membrane fragments were extracted using the membrane and cytosol protein extraction kit from Beyotime (P0033). Hepa1-6 cells were grown in cell culture dish to 80%-90% density and then detached by cell scrapers. The cells were washed with PBS for three times by centrifugation at 2,000 rpm for 5 min. The collected cells were suspended in membrane protein extraction buffer solutions containing phenylmethanesulfonyl fluoride (PMSF, 1 mM) and incubated in ice bath for 10 min. Subsequently, the cells were broken by the Ultrasonic Homogenizer (40 W, 2 min). Then, the mixture was centrifuged at 3,000 rpm for 10 min at 4 °C and the collected suspension was further centrifuged at 15,000 rpm for 60 min at 4 °C to obtain Hepa1-6 cell membrane fragments. Finally, the Hepa1-6 cell membrane was resuspended in PBS, and stored at -80 °C.

**Preparation of NMOF-Fe/Cu-Dox@M:** Hepa1-6 cell membranes were extruded through polycarbonate porous membranes (pore diameters: 1000 nm) for 7 times and the uniform size vesicles were obtained. Then, the vesicles were mixed with NMOF-Fe/Cu-Dox@M obtained previously (mass ratio = 1:2) and extruded over 7 times through microporous membrane (pore diameter: 1000 nm). At last, the NMOF-Fe/Cu-Dox@M was obtained and stored at 4 °C for further experiments.

**TMB Catalytic Reaction:** 100 mg TMB was added to 10 mL DMSO solvent to obtain a 10 mg·mL$^{-1}$ TMB solution. The above solution (1 mL) was transferred into a centrifuge tube with the addition of 9 mL NaAc/HAc buffer (pH 5, 0.1 mM). Then, 100 μL NMOF-Fe/Cu (100 μg·min$^{-1}$), 200 μL TMB (1 mg·min$^{-1}$) solution, and $H_2O_2$ (10 mM) were mixed, and diluted to 1 mL with NaAc/HAc buffer. The solutions were subjected to UV measurement at 652 nm after incubation for different time durations.

**POD Like Catalytic Activity of NMOF-Fe/Cu:** Generation of •OH was studied by MB and TA assay. Briefly, MB aqueous solution (5 μg·min$^{-1}$), $H_2O_2$ (10 mM) and NMOF-Fe/Cu (100 μg·mL$^{-1}$) were mixed and incubated for 12 h. The UV–vis absorption spectra were measured. Similarly, TA aqueous solution (0.5 mM), $H_2O_2$ (100 mM), and NMOF-Fe/Cu (100 μg·mL$^{-1}$) were mixed and incubated for 2 h. The fluorescence emission spectra of the TA were recorded at 425 nm.

**Kinetic Measurements:** Kinetic measurements were carried out with different concentrations of $H_2O_2$ (0, 25, 100, 200, 400, 1000 μM). The solutions were subjected to UV measurement at 652 nm after incubation for different time durations. The $K_m$ and $V_m$ constants were obtained by fitting Michaelis–Menten equation. The equation is $V = V_{max} \times C/(K_m + C)$, where V is the initial catalyzed reaction rate; C is the initial concentration of TMB; $V_{max}$ points the maximal catalyzed reaction rate; $K_m$ represents to the Michaelis constant.

**Detection of GSH:** The consumption of GSH was measured using DTNB as a probe. Firstly, DTNB (10 mM) was used to detect the absorption at 412 nm of GSH (5 mM). The GSH consumption assay of different materials were divided the experiment into four groups: Group 1, GSH; Group 2, GSH+NMOF-Fe/Cu; Group 3, GSH+NMOF-Fe;

Group 4, GSH+NMOF-Cu. After the four experimental groups reacted in Tris-HCl (pH 8.0) for 30 min, the solution was centrifuged and the supernatant was mixed with DTNB and the absorption at 412 nm was detected by a microplate reader. Next, the concentration of GSH after reaction with NMOF-Fe/Cu in the same conditions was detected according to the above steps.

***In vitro* DOX Release:** The standard fluorescence curve of Dox was firstly plotted and loading efficiency of Dox by NMOF-Fe/Cu-Dox was then calculated. 1 mg of NMOF-Fe/Cu-Dox powder was dispersed in 10 mL of PBS at different pH values in 15 mL tubes and shaken at 37 °C. At different time points, the supernatants are collected and centrifugated. The Dox concentration in the supernatant solution was measured by fluorescence spectroscopy ($\lambda_{ex}$ = 495 nm, $\lambda_{exm}$ = 595 nm). The loading efficiency is calculated as follows: Dox encapsulation efficiency = (initial Dox - supernatant Dox)/initial Dox. The pH-dependent release of Dox from NMOF-Fe/Cu-Dox was investigated in PBS at different pH values (6.5 and 7.4).

**Cell Culture:** Hepa1-6 cells, HUVEC cells and NIH3T3 cells were grown in high glucose DMEM supplemented with 10% FBS and 1% penicillin-streptomycin at 37 °C with 5% $CO_2$.

**Cytotoxicity assay:** The cytotoxicity of different treatments to Hepa1-6 cells, HUVEC cells and NIH3T3 cell were tested by CCK-8 assay (Sihai Bio-Tech Co., Ltd, Shanghai). $5\times10^3$ of Hepa1-6 cells were cultured for 24 h in 96-well plates. To simulate the acidic microenvironment in the solid tumor, DMEM (pH 7.4) was acidized to pH 6.5 by hydrochloric acid. Then, different nanoparticles (NMOF, NMOF-Fe/Cu, NMOF-

Fe/Cu-Dox, NMOF-Fe/Cu-Dox@M) at various concentrations (10 μg·mL$^{-1}$, 20 μg·mL$^{-1}$, 40 μg·mL$^{-1}$, 60 μg·mL$^{-1}$, 80 μg·mL$^{-1}$) were added to the cells and cultured for another 12 h or 24 h. Finally, the cells were cultured with CCK-8 solution (10% in DMEM) for 1 h, the cell proliferation was determined using the microplate reader by comparing the absorbance at 450 nm to the control group.

**Flow cytometry was conducted for cell apoptosis analysis：** Hepa1-6 cells were seeded ($3\times10^5$ cells) in 6-well plates and cultured for 24 h. Then the culture media were replaced by DMEM containing NMOF, NMOF-Fe/Cu, NMOF-Fe/Cu-Dox and NMOF-Fe/Cu-Dox@M (60 μg·mL$^{-1}$) and incubated for another 12 h. After co-incubation, the cells were collected by centrifugation. Then, the Annexin V-FITC/PI Apoptosis Detection Kit was used to stain the cells for 10 min before the flow cytometry analysis.

***In vitro* Live/Dead Cell Staining:** *In vitro* live/dead cell staining was performed with Calcein-AM (for live cells) and propidiumiodide (PI, for dead cells). $1\times10^5$ of Hepa1-6 cells were seeded into 48 well plates and cultured for 24 h. The cells were then incubated with fresh mediums containing NMOF, NMOF-Fe/Cu, NMOF-Fe/Cu-Dox and NMOF-Fe/Cu-Dox@M (60 μg·mL$^{-1}$) for 12 h, respectively. Then, the culture media were replaced by 200 μL DMEM containing 0.5 μM Calcein-AM and 2 μM PI. After incubation for another 30 min, the cells were washed with PBS for three times. Afterwards, the influence of concentration on the live/dead cell staining at various concentrations NMOF-Fe/Cu-Dox@M (10 μg·mL$^{-1}$, 20 μg·mL$^{-1}$, 40 μg·mL$^{-1}$, 60 μg·mL$^{-1}$, 80 μg·mL$^{-1}$) was studied. Hepa1-6 cells were seeded ($1\times10^5$ cells) in the 48-well plates and cultured for 12 h. Then, the culture media were replaced by 200 μL DMEM containing 0.5 μM Calcein-AM and 2 μM PI and incubated for 30 min. The cells were washed with PBS three times and the fluorescence imaging was acquired by an inverted fluorescence microscope.

**Intracellular ROS Evaluation:** Hepa1-6 cells were seeded ($1\times10^5$ cells) in the 48-well plates. The culture media were then replaced by DMEM containing NMOF, NMOF-Fe/Cu, NMOF-Fe/Cu-Dox and NMOF-Fe/Cu-Dox@M (40 $\mu g\cdot mL^{-1}$). After incubation for 4 h, the culture media were replaced by 200 μL DCFH-DA (10 μM in FBS-free DMEM) and incubated for 30 min. The cells were washed with PBS three times and the level of intracellular ROS was evaluated by detecting the fluorescence ($\lambda_{ex}$ = 488 nm, $\lambda_{em}$ = 525 nm). Then, Hepa1-6 cells were seeded ($1\times10^5$ cells) in the 48-well plates and cultured for 12 h with different dosages of NMOF-Fe/Cu-Dox@M (10 $\mu g\cdot mL^{-1}$, 20 $\mu g\cdot mL^{-1}$, 40 $\mu g\cdot mL^{-1}$, 60 $\mu g\cdot mL^{-1}$, 80 $\mu g\cdot mL^{-1}$). After incubation for 4 h, the culture media were replaced by 200 μL DCFH-DA (10 μM in FBS-free DMEM) and incubated for 30 min. The cells were washed with PBS three times and the intracellular ROS observed under a fluorescent inverted microscope.

**Intracellular Uptake of NMOF-Fe/Cu-Dox@M:** The uptake of NMOF-Fe/Cu-Dox@M and NMOF-Fe/Cu-Dox were detected through the fluorescence of Dox with the help of confocal laser scanning microscope (CLSM). In detail, $3\times10^5$ of different cells (Hepa1-6 cells, HUVEC cells and NIH3T3 cells) were seeded in 24 well-plate and cultured for 24 h. Then the cells were incubated with NMOF-Fe/Cu-Dox@M and NMOF-Fe/Cu-Dox for 1 h, 4 h or 8 h. After being washed with PBS, fixed with 4% paraformaldehyde, and then dyed with DAPI (2 $\mu g\cdot mL^{-1}$, 10 min), the cells were used for confocal cell imaging by CLSM.

**Mitochondria and Lysosomes Staining:** In detail, 3×10$^5$ Hepa1-6 cells were seeded in 24 well-plate and cultured for 24 h. Then the Cy5 labeled NMOF-Fe/Cu were mixed with Hepa1-6 cell membranes obtained previously (mass ratio = 1:2) and extruded over 7 times through microporous membrane (pore diameter: 1000 nm). The Cy5 labeled NMOF-Fe/Cu@M (40 μg·mL$^{-1}$) was added and incubated for 4 h. Next, the cells were washed with PBS three times and stained with Mito-Tracker Green (1 μM, 15 min) and Lyso-Tracker Green (200 nM, 15 min). Then fixed with 4% paraformaldehyde, anddyed with DAPI (2 μg·mL$^{-1}$, 10 min), the cells were used for confocal cell imaging by CLSM.

**Detection of LPO:** The LPO was detected by $C_{11}$-BODIPY™ 581/591 (Invitrogen™). The cells were incubated with Cy5 labeled NMOF-Fe/Cu@M (80 μg·mL$^{-1}$) for 6 h. After being washed with PBS three times and incubated with C11-BODIPY™ 581/591 (5 μM, 15 min). Then fixed with 4% paraformaldehyde, and dyed with DAPI (2 μg·mL$^{-1}$, 10 min), the cells were used for confocal cell imaging by CLSM.

**Detection of JC-1:** Mitochondrial membrane potential was measured using JC-1 (MedChemExpress, CBIC2). In normal conditions, JC-1 is accumulated in the matrix of mitochondria and forms J-aggregates ($\lambda_{ex}$ = 585 nm; $\lambda_{em}$ = 590 nm), which produces red fluorescence. When the mitochondrial membrane potential is lost, JC-1 forms a monomer and exists in the cytosol ($\lambda_{ex}$ = 514 nm; $\lambda_{em}$ = 529 nm) that produces green fluorescence. The cells were incubated with Cy5 labeled NMOF-Fe/Cu@M for 6 h. After being washed with PBS three times and incubated with JC-1 (5 μg·mL$^{-1}$,15 min). Then fixed with 4% paraformaldehyde, and then dyed with DAPI (2 μg·mL$^{-1}$, 10 min), the cells were used for confocal cell imaging by CLSM.

**SDS-PAGE:** The total protein distributions of Hepa1-6 cell membrane and NMOF-Fe/Cu-Dox@M were visualized by SDS-PAGE. In brief, the samples were lysed in radioimmunoprecipitation assay (RIPA) buffer and quantified by BCA protein assay. Then the solutions were mixed with loading buffer and heated at 100 °C for 10 min. The solutions with same amounts of protein were loaded into 12% polyacrylamide gels and the electrophoresis was processed at 80 mV. Finally, the gel was immersed in Coomassie brilliant blue buffer for 12 h and then shook slightly in destaining solution for 2 days.

**Proteomic and Bioinformatic Analyses:** Protein in the supernatant was precipitated with TCA precipitation assay and the precipitation was resuspended in redissolved solution (8 M Urea/100 mM Tris-HCl, pH = 8.5). After the protein precipitation was redissolved, the protein concentration was determined by BCA method. With the same amount of protein among different samples, the redissolved solution was used to fill all samples to the same volume. Protein reduction and alkylation were conducted with TCEP and CAA at 37°C for 1 h. Urea was diluted below 2 M using 100 mM Tris-HCl. Trypsin was added at a ratio of 1:50 (enzyme: protein, w/w) for overnight digestion at 37°C. The next day, TFA was used to bring the pH down to 6.0 to end the digestion. After centrifugation (12000 g, 15 min), the supernatant was subjected to peptide purification using a self-made SDB-RPS desalting column. The peptide eluate was vacuum dried and stored at -20 °C for later use. All samples were analyzed on Q Exactive HF mass spectrometer (Thermo, Fisher). An UltiMate 3000 RSLCnano system (Thermo) was coupled to Q Exactive HF with a CaptiveSpray nano ion source

(Bruker Daltonics). Peptide samples were injected into a C18 Trap column (75 μm*25cm, 1,9 μm particle size, 120 Å pore size, Thermo), and separated in a reversed-phase C18 analytical column (75 μm*15 cm, 1.7 μm particle size, 100 Å pore size, Ion Opticks). Mobile phase A (0.1% formic acid in water) and mobile phase B (3% DMSO in 80% ACN) were used to establish the separation gradient at a flow rate of 300 nL/min. The MS data acquisition was performed in DDA mode. The MS and MS/MS spectra were acquired from 350 to 1500 m/z. MS raw data were analyzed with MaxQuant (V2.0.1.0) using the Andromeda database search algorithm. At last, both proteins presented in NMOF-Fe/Cu-Dox@M and Hepa1-6 cells were considered as membrane-reserved proteins. The bioinformatic analyses of membrane proteins were performed by the online software Database (https://www.bioinformatics.com.cn/).

**DLAT immunofluorescence:** $3\times10^5$ cells were seeded on coverslips in 24-well plate overnight and treated as indicated. Briefly, the cells were fixed with 4% paraformaldehyde for 15 min, the cells were washed three times with PBS. The cells were permeabilized in 0.5% Triton X-100 for 10 min and blotted with 3% BSA for 1 h. The cells were incubated with DLAT antibody (1:3000) at 4 °C overnight. Then, the cells were washed three times with PBS, incubated with the appropriate secondary antibodies for 1 h at 37°C, washed and sealed with mounting medium including DAPI (2 $ug\cdot mL^{-1}$). Images were captured on confocal.

**Measurement of Intracellular GSH levels:** Hepa1-6 cells in 24-well plates ($3\times 10^5$ cells per well) were grown overnight to reach 70-90% confluence, followed by incubation with NMOF-Fe/Cu-Dox@M (80 μg·mL$^{-1}$) for 12 h. After washing three times with PBS, the amounts of GSH were detected using GSH and GSSG assay kits according to the manufacturer's instructions.

**RNA-Seq analysis:** Total RNA was extracted using Nucleozol (Gene, 7040404). The concentrations of total RNA samples were determined by a Nanodrop 2000C (Thermo Scientific, USA) their RNA integrity was assessed by an Agilent 2100 Bioanalyze. The RNA-Seq was performed on Illumina Novaseq 6000 platform with service provided by Anoroad Co. Ltd. (Beijing, China).

**In vivo anti-tumor therapy/in vivo anti-tumor immunotherapy**: Male C57/B6L mice aged 5-6 weeks (20-22 g) were purchased from Beijing Vital River Laboratory Animal Technology Co., Ltd. All animal experiments in this study were approved by the Animal Use Committee of Tongji Hospital, Tongji Medical College, Huazhong University of Science and Technology. Hepa1-6 ($2\times10^6$) cells were subcutaneously inoculated into the right flanks of C57BL/6 mice. C57/B6L mice bearing Hepa1-6 tumors were assigned into five groups at random ($n$=5 for each group) as follows: PBS (control), NMOF-Fe/Cu, NMOF-Fe/Cu-Dox and NMOF-Fe/Cu-Dox@M. Different formulations of nanoparticles were intravenously injected into the mice at a dose of 200 μg (2 mg/mL) every 2 days for three times. The intraperitoneal injection of αPD-1 at a dose of 100 μg (1 mg/mL) every 3 days for three times. At the end of the experiments, the mice were sacrificed, and the tumors were collected, weighed and analyzed by Ki67, H&E, and TUNEL staining. The major organs, including the heart, liver, spleen, lung, and kidney, were also harvested and analyzed by H&E staining.

**In vivo tumor biodistribution of NMOF-Fe/Cu-Dox@M**: Hepa1-6 ($2\times10^6$) cells were subcutaneously injected into Male C57/B6L mice. When the tumor volume reached ≈100 $mm^3$, mice were randomly assigned into two different groups group to evaluate the biodistribution of NMOF-Fe/Cu-Dox@M. Dir-labeled NMOF-Fe/Cu-Dox and NMOF-Fe/Cu-Dox@M were injected into mice via the tail vein. The tumor region was imaged at the indicated times (2, 4, 8, 12, and 24 h). Moreover, the tumor-bearing mice were sacrificed, and major organs were obtained and imaged.

**Western Blot Assay**: After different formulations treated, cells were collected and harvested using RIPA Lysis Buffer to obtained proteins. Use the BCA protein detection kit (Beyotime Co., Ltd.) for protein quantification and dilute it to the same concentration. These proteins were then separated by SDS-PAGE gradient gel and transferred to PVDF membrane. Then, the membranes were incubated with different antibodies and β-actin at 4 °C overnight. The protein bands were performed using the enhanced chemiluminescence solution reaction. We have provided uncropped and unprocessed scans of the most important blots in the source data file.

**Hemolysis Assay**: Fresh blood (1 mL) was washed with PBS (10 mL) three times (1500 rpm, 15 min, 4 °C) and resuspended with PBS to obtain 2% red cell suspension. The normal saline and the deionized water were the negative and positive control groups, respectively. After incubation at 37 °C for 4 h, the mixture was then centrifuged. The absorption of supernatant was measured by a BioTek NEO2 microplate reader at 577 nm. Each group contained three parallel samples. Hemolysis rate (%) = ($OD_{test}$ - $OD_{negative\ control}$)/($OD_{positive\ control}$ - $OD_{negative\ control}$) × 100%

**Statistical Analysis**: All the data were presented as mean ± standard deviation. Differences among samples were calculated with the two-tailed Student's t-test using an independent samples test in Origin. Differences among groups were considered statistically significant at $p < 0.05$ (*$p < 0.05$, ** $p < 0.01$, *** $p < 0.001$ and ****$p < 0.0001$).

### Supplementary figures

**Supplementary Fig. S1 (a) The SEM image of NMOF-Fe/Cu; (b) NMOF-Fe/Cu-Dox.**

**Supplementary Fig. S2 The SEM and TEM image of NMOF-Fe/Cu-Dox@M.**

**Supplementary Fig. S3 TGA curve of both NMOF and NMOF-Fe/Cu NMOF and NMOF-Fe/Cu under $N_2$ atmosphere.**

**Supplementary Fig. S4 FTIR spectra of NMOF and NMOF-Fe/Cu.**

**Supplementary Fig. S5 XPS broad spectra of NMOF and NMOF-Fe/Cu.**

**Supplementary Fig. S6 XRD of NMOF, NMOF-Fe/Cu and NMOF-Fe/Cu-Dox.**

**Supplementary Fig. S7 Hydrodynamic size of NMOF, NMOF-Fe/Cu, NMOF-Fe/Cu-Dox and NMOF-Fe/Cu-Dox@M.**

**Supplementary Fig. S8 (a) The hydrodynamic size; (b) the *Zeta* potentials of**

**NMOF-Fe/Cu-Dox@M in water and 10% FBS for 1, 3, 5 and 7 days.**

**Supplementary Fig. S9 SDS-PAGE protein analysis of markers, NMOF-Fe/Cu-Dox@M and Heap 1-6 cells membrane.**

**Supplementary Fig. S10 Probing •OH formation through the fluorescence spectrum of the adduct of •OH with TA.**

**Supplementary Fig. S11 Absorption spectra corresponding to the oxidized TMB ($TMB_{ox}$) generated by different ratios of Cu and Fe in NMOF-Fe/Cu.**

**Supplementary Fig. S12 Absorption spectra corresponding to the oxidized TMB ($TMB_{ox}$) generated by NMOF-Fe/Cu at different time.**

**Supplementary Fig. S13 The absorbance spectrum of GSH treated with different materials (30 min).**

**Supplementary Fig. S14 The non-linear regression curve of Dox.**

**Supplementary Fig. S15 (a), (b) Confocal images of the uptake of the NMOF-Fe/Cu-Dox and NMOF-Fe/Cu-Dox@M by HUVEC cells at 1 h, 4 h and 8 h, respectively. Scale bar: 20 μm.**

**Supplementary Fig. S16 Relative fluorescence intensities of HUVEC cells incubated with NMOF-Fe/Cu-Dox and NMOF-Fe/Cu-Dox@M for 1 h, 4 h and 8 h (*n*=3).**

**Supplementary Fig. S17 Fluorescence image of subcellular location of Cy5-labled NMOF-Fe/Cu-Dox@M after 4 h. Cy5 fluorescence (red), Mito Tracker Green (detect mitochondria) and DAPI (blue). Pearson's correlation coefficient and plot profile analysis of Mito Tracker co-localization with NMOF-Fe/Cu-Cy5@M. The red lines representing NMOF-Fe/Cu-Cy5@M showed trajectories that were roughly similar to the black lines representing Mito tracker Green, indicating significant co-localization.**

**Supplementary Fig. S18 (a) Cell viability of HUVEC cells after incubated with NMOF-Fe/Cu-Dox@M for 12 h (*n*=3); (b) Cell viability of NIH3T3 cells after incubated with NMOF-Fe/Cu-Dox@M for 12 h (*n*=4).**

**Supplementary Fig. S19 Cell viability of Hepa1-6 cells after incubated with NMOF-Cu, NMOF-Fe, NMOF-Fe/Cu for 12 h (*n*=3).**

**Supplementary Fig. S20 Cell viability of Hepa1-6 cells after incubated with NMOF,**

**NMOF-Fe/Cu, NMOF-Fe/Cu-Dox and NMOF-Fe/Cu-Dox@M for 4 h, respectively (*n*=3).**

**Supplementary Fig. S21 Cell viability of Hepa1-6 cells after incubated with NMOF, NMOF-Fe/Cu, NMOF-Fe/Cu-Dox and NMOF-Fe/Cu-Dox@M for 12 h in pH 6.5 (*n*=3).**

**Supplementary Fig. S22 The live/dead staining of Hepa1-6 cells after the NMOF, NMOF-Fe/Cu, NMOF-Fe/Cu-Dox and NMOF-Fe/Cu-Dox@M. scale bar: 20 μm.**

**Supplementary Fig. S23 The live/dead stained images of Hepa1-6 treated with different concentrations of NMOF-Fe/Cu-Dox@M. scale bar: 20 μm.**

**Supplementary Fig. S24 ROS production was detected by DCFH-DA in Hepa1-6 cells at different concentrations. scale bar: 20 μm.**

**Supplementary Fig. S25 Western blot analysis of GPX4, Cleaved- parp1, ACSL4, FDX1 expression after incubated with different concentrations of NMOF-Cu/Fe-DOX@M.**

**Supplementary Fig. S26 The results of the hemolysis experiment of NMOF-Fe/Cu-Dox@M (*n*=3).**

**Supplementary Fig. S27 Bioluminescence images of major organs of mice after tail vein injection of MOF-Fe/Cu-Dir@M for 12 h.**

**Supplementary Fig. S28 The fluorescence intensity of various group of tumors in Day 0 and Day12 (*n*=5).**

**Supplementary Fig. S29 H&E histopathological analysis of major organ tissues after various treatments.**

**Supplementary Fig. S30 Flow gating strategy of myeloid cells in the tumor.**

**Supplementary Fig. S31 Representative flow cytometric plots of M2 macrophages and quantification of M2 macrophages after different treatments (*n*=5). Statistical significance was calculated by *t*-test.**

**Supplementary Fig. S32 Flow gating strategy of (a) macrophages and (b) CD8+ T cells.**

**Supplementary Fig. S33 Percentages of Granzyme $B^+$, and PD-$1^+$， Ki$67^+$ and Tim$3^+$ T cells in $CD8^+$ T cells extracted from the spleen cultured with RAW264.7 cells after co-culturing. (1) CM, (2) CM Hepa1-6 and (3) CD3/CD28 -treated**

**groups, respectively (*n*=3).**

**Supplementary Fig. S34 The cytotoxic effects of co-cultured $CD8^+$ T cells on Hepa1-6 cells, (1) CM, (2) CM Hepa1-6 and (3) CD3/CD28-treated groups, respectively (*n*=5).**

**Supplementary Fig. S35 The fluorescence intensity of various group of tumors in Day0 and Day12. (1) PBS; (2) NMOF-Fe/Cu-Dox@M; (3) αPD-1; (4) NMOF-Fe/Cu-Dox@M+αPD-1 (*n*=5).**

**Supplementary Fig. S36 Tumor tissue staining by H&E, Ki67, TUNEL after antitumor treatment and immunotherapy.**

**Supplementary Fig. S37 (a)The fluorescence intensity of Ki67 after various treatments; (b) The fluorescence intensity of Tunel after various treatments. (1) PBS; (2) NMOF-Fe/Cu-Dox@M; (3) αPD-1; (4) NMOF-Fe/Cu-Dox@M+αPD-1 (*n*=5).**

**Supplementary Fig. S38 The body weight variation in different treatments (*n*=5).**

**Supplementary Fig. S39 H&E histopathological analysis of major organ tissues after various treatments.**

**Supplementary Fig. S40 The indices of liver function, renal function indicator and blood cell index in mouse serum after 12-day treatment. (1) PBS; (2) NMOF-Fe/Cu-Dox@M; (3) αPD-1; (4) NMOF-Fe/Cu-Dox@M+αPD-1 (*n*=5).**

**Supplementary Fig. S41 Flow gating strategy of mouse immune T cells.**

**Supplementary Fig. S42 The quantification of $CD8^+$ T cells (a), Treg cells (b), $CD4^+$/$IFN\gamma^+$ T cells (c), $CD4^+$ Tcm cells (d) surface marker expression in tumors of mice after different treatments (*n*=5). (1) PBS; (2) NMOF-Fe/Cu-Dox@M; (3) αPD-1; (4) NMOF-Fe/Cu-Dox@M+αPD-1.**

### Supplementary tables

**Supplementary table 1. The concentration of metal ion.**

**Supplementary table 2. Antibodies used for this work.**

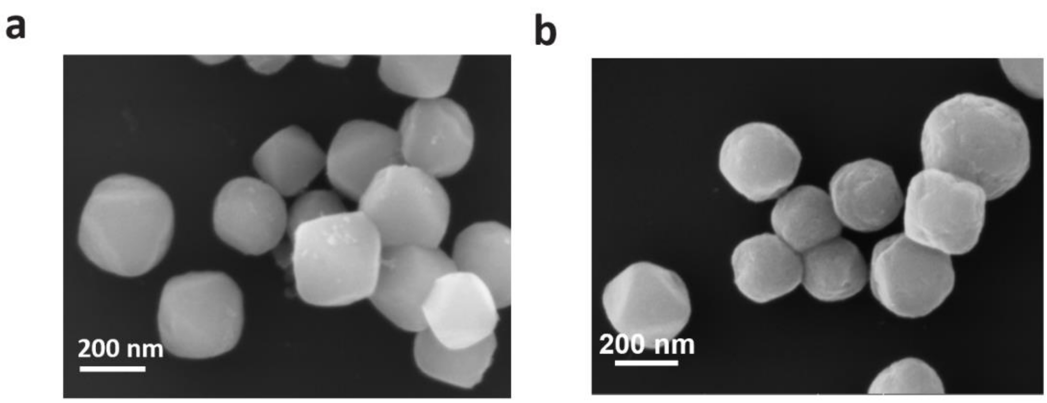

**Supplementary Fig S1** (a) The SEM image of NMOF-Fe/Cu; (b) NMOF-Fe/Cu-Dox.

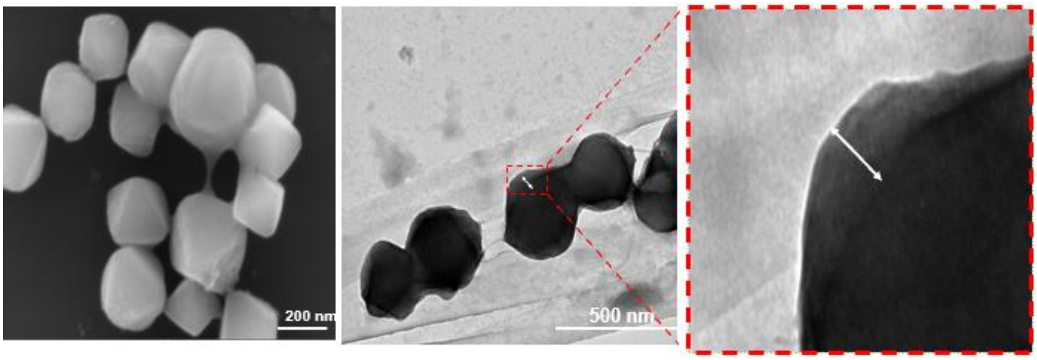

**Supplementary Fig. S2** The SEM and TEM image of NMOF-Fe/Cu-Dox@M.

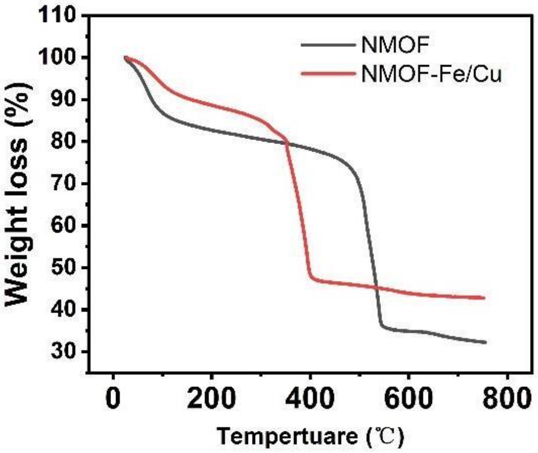

**Supplementary Fig. S3** TGA curve of both NMOF and NMOF-Fe/Cu NMOF and NMOF-Fe/Cu under $N_2$ atmosphere.

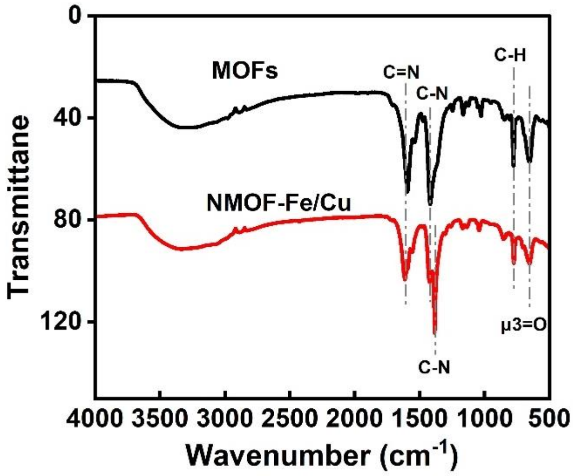

**Supplementary Fig. S4** FTIR spectra of NMOF and NMOF-Fe/Cu.

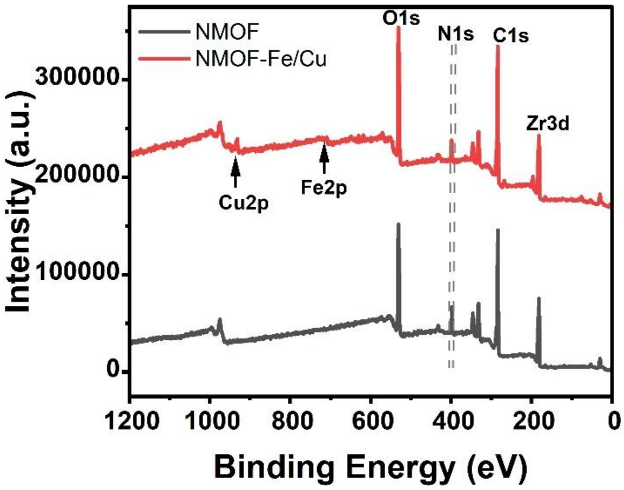

**Supplementary Fig. S5** XPS broad spectra of NMOF and NMOF-Fe/Cu.

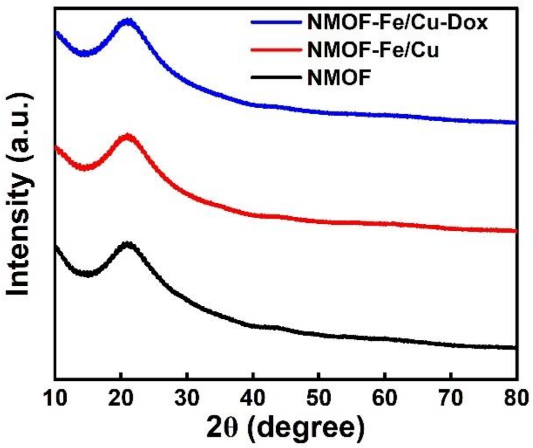

**Supplementary Fig. S6** XRD of NMOF, NMOF-Fe/Cu and NMOF-Fe/Cu-Dox.

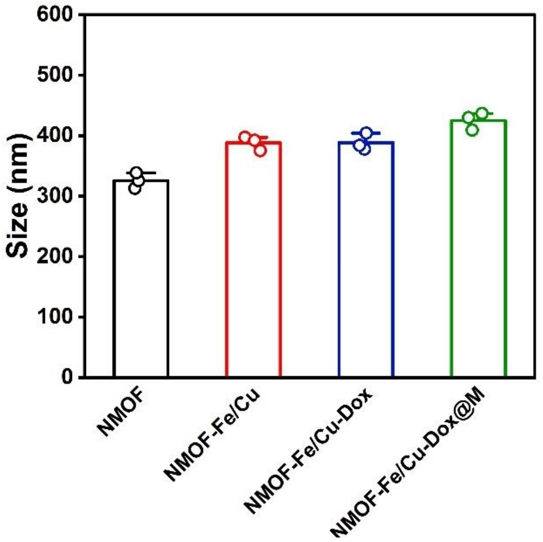

**Supplementary Fig. S7** Hydrodynamic size of NMOF, NMOF-Fe/Cu, NMOF-Fe/Cu-Dox and NMOF-Fe/Cu-Dox@M.

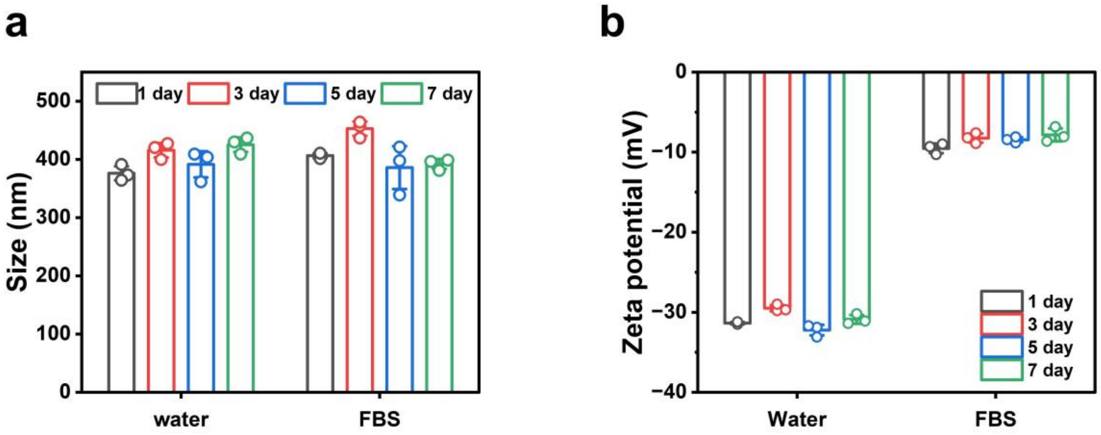

**Supplementary Fig. S8** (a) The hydrodynamic size; (b) the *Zeta* potentials of NMOF-Fe/Cu-Dox@M in water and 10% FBS for 1, 3, 5 and 7 days.

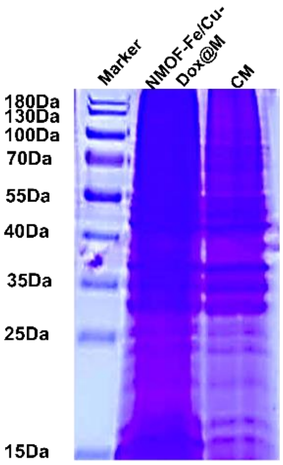

**Supplementary Fig. S9** SDS-PAGE protein analysis of markers, NMOF-Fe/Cu-Dox@M and Heap 1-6 cells membrane.

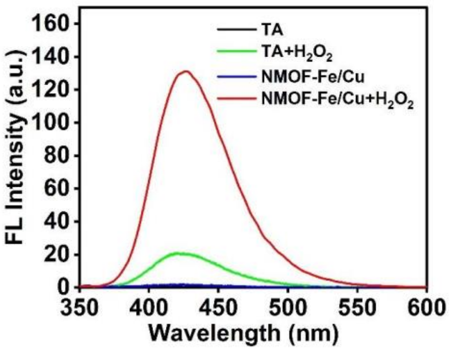

**Supplementary Fig. S10** Probing •OH formation through the fluorescence spectrum of the adduct of •OH with TA.

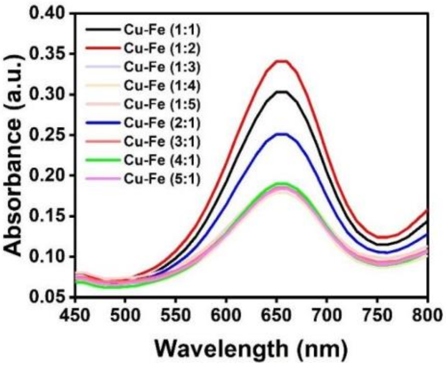

**Supplementary Fig. S11** Absorption spectra corresponding to the oxidized TMB ($TMB_{ox}$) generated by different ratios of Cu and Fe in NMOF-Fe/Cu.

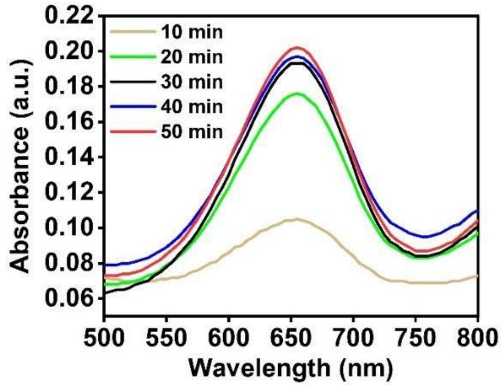

**Supplementary Fig. S12** Absorption spectra corresponding to the oxidized TMB ($TMB_{ox}$) generated by NMOF-Fe/Cu at different time.

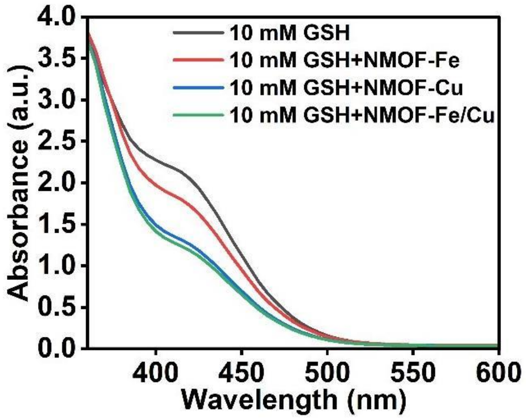

**Supplementary Fig. S13** The absorbance spectrum of GSH treated with different materials (30 min).

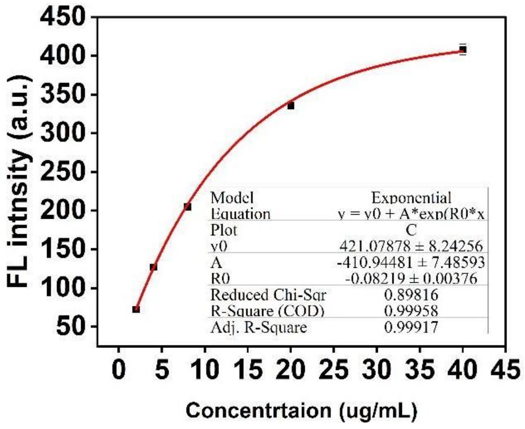

**Supplementary Fig. S14** The non-linear regression curve of Dox.

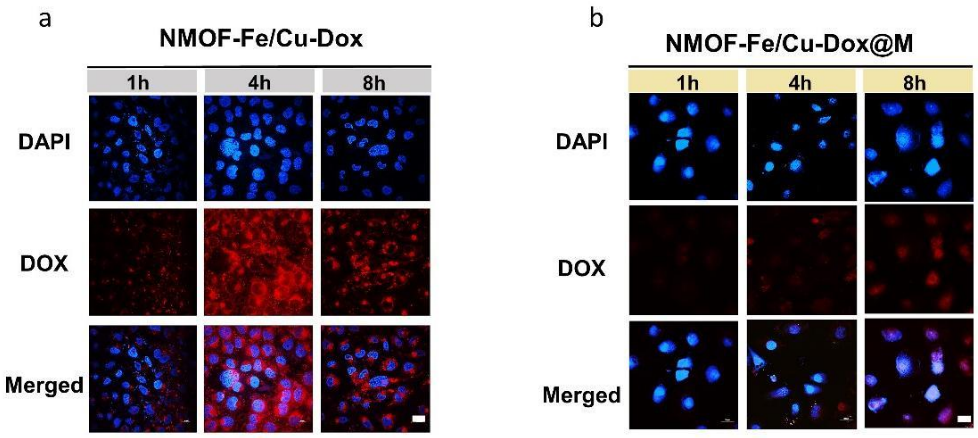

**Supplementary Fig. S15** (a), (b) Confocal images of the uptake of the NMOF-Fe/Cu-Dox and NMOF-Fe/Cu-Dox@M by HUVEC cells at 1 h, 4 h and 8 h, respectively. Scale bar: 20 μm.

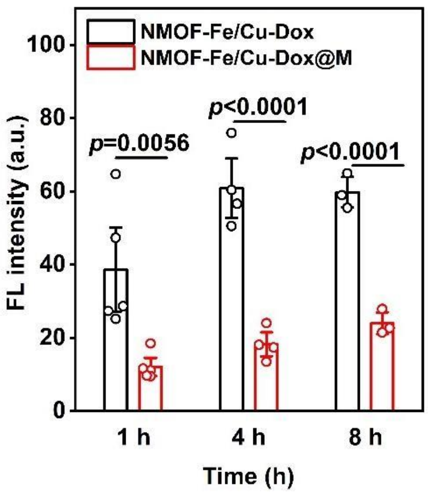

**Supplementary Fig. S16** Relative fluorescence intensities of HUVEC cells incubated with NMOF-Fe/Cu-Dox and NMOF-Fe/Cu-Dox@M for 1 h, 4 h and 8 h ($n$=3).

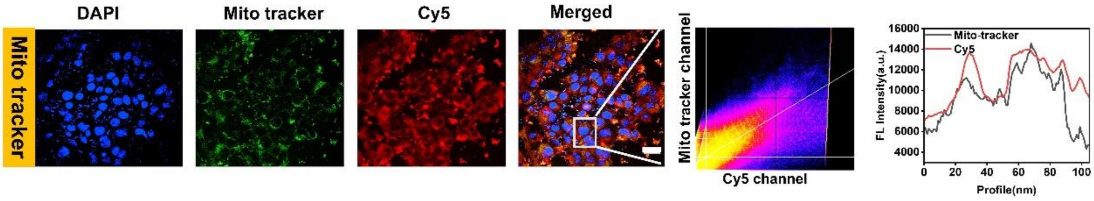

**Supplementary Fig. S17** Fluorescence image of subcellular location of Cy5-labled NMOF-Fe/Cu-Dox@M after 4 h. Cy5 fluorescence (red), Mito Tracker Green (detect mitochondria) and DAPI (blue). Pearson's correlation coefficient and plot profile analysis of Mito Tracker co-localization with NMOF-Fe/Cu-Cy5@M. The red lines representing NMOF-Fe/Cu-Cy5@M showed trajectories that were roughly similar to the black lines representing Mito tracker Green, indicating significant co-localization.

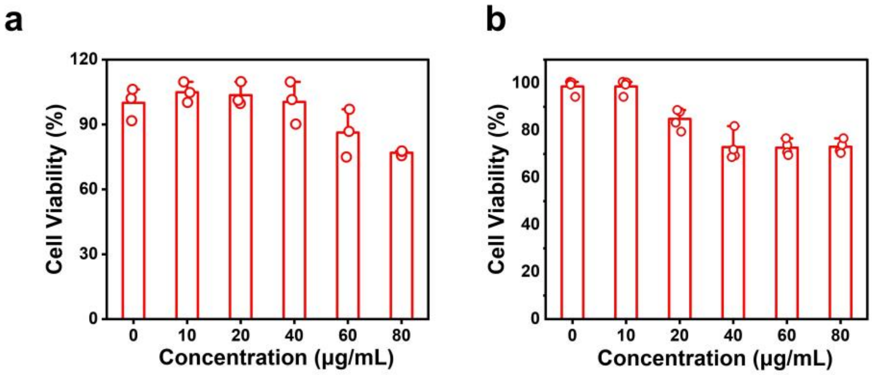

**Supplementary Fig. S18** (a) Cell viability of HUVEC cells after incubated with NMOF-Fe/Cu-Dox@M for 12 h ($n$=3); (b) Cell viability of NIH3T3 cells after incubated with NMOF-Fe/Cu-Dox@M for 12 h ($n$=4).

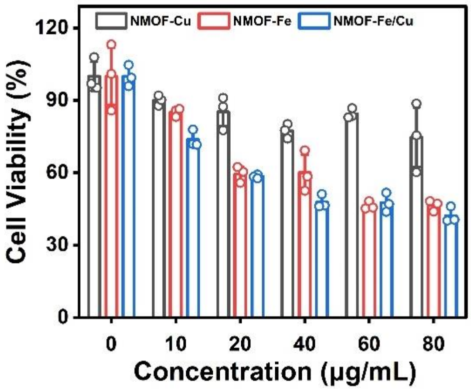

**Supplementary Fig. S19** Cell viability of Hepa1-6 cells after incubated with NMOF-Cu, NMOF-Fe, NMOF-Fe/Cu for 12 ($n$=3).

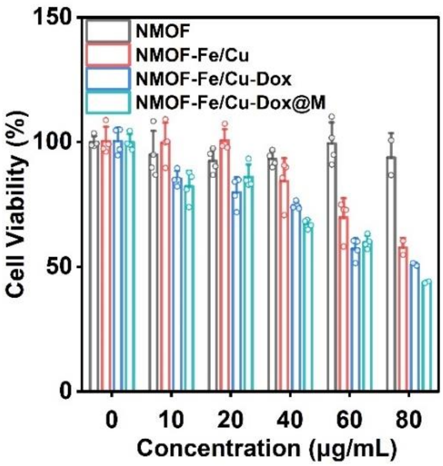

**Supplementary Fig. S20** Cell viability of Hepa1-6 cells after incubated with NMOF, NMOF-Fe/Cu, NMOF-Fe/Cu-Dox and NMOF-Fe/Cu-Dox@M for 4 h, respectively (*n*=3).

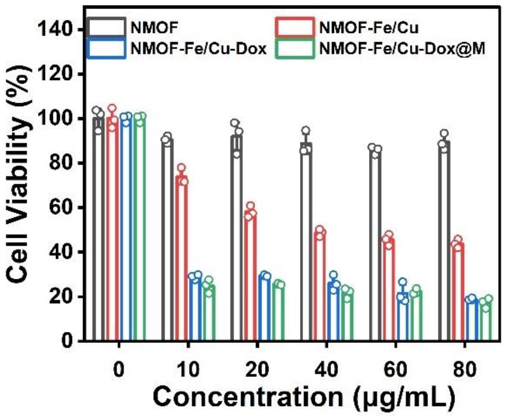

**Supplementary Fig. S21** Cell viability of Hepa1-6 cells after incubated with NMOF, NMOF-Fe/Cu, NMOF-Fe/Cu-Dox and NMOF-Fe/Cu-Dox@M for 12 h in pH 6.5 (*n*=3).

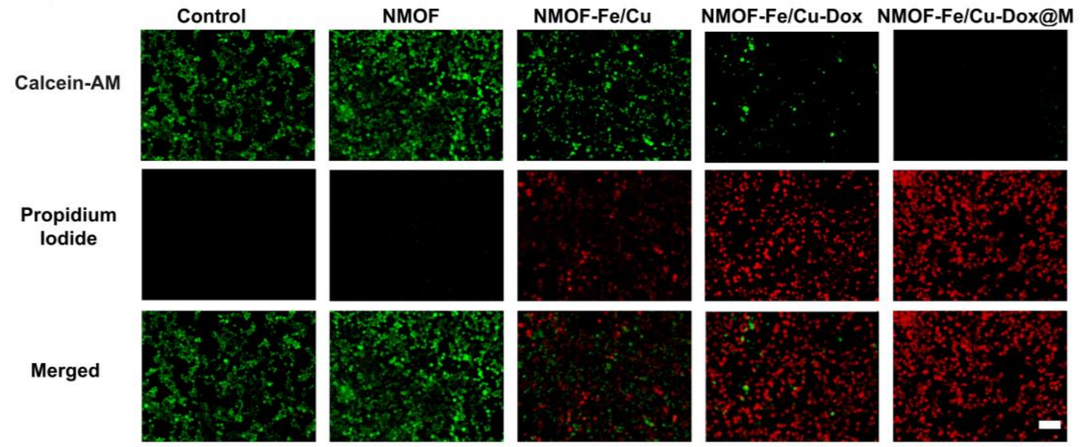

**Supplementary Fig. S22** The live/dead staining of Hepa1-6 cells after the NMOF, NMOF-Fe/Cu, NMOF-Fe/Cu-Dox and NMOF-Fe/Cu-Dox@M treatments. scale bar: 20 μm.

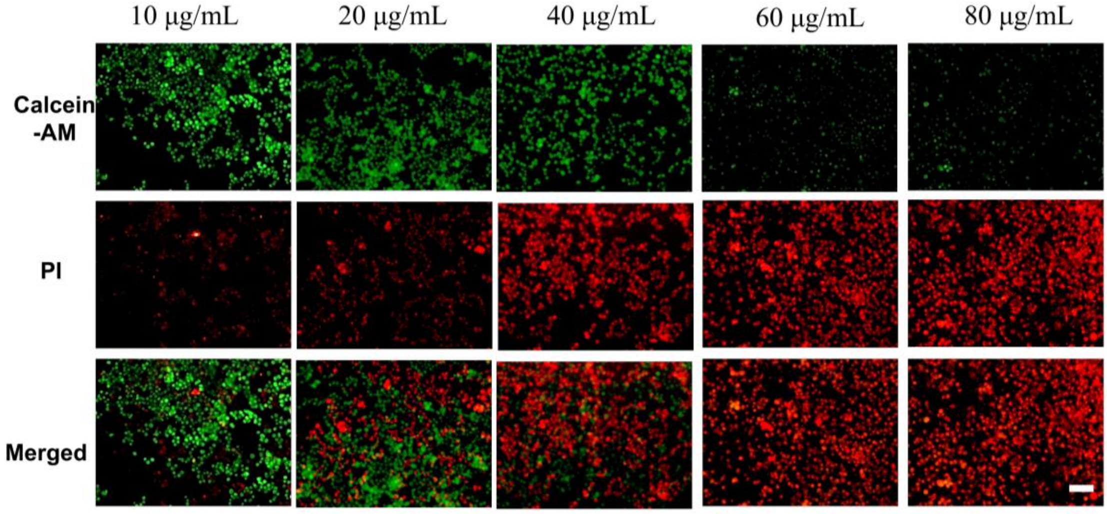

**Supplementary Fig. S23** The live/dead stained images of Hepa1-6 treated with different concentrations of NMOF-Fe/Cu-Dox@M. scale bar: 20 μm.

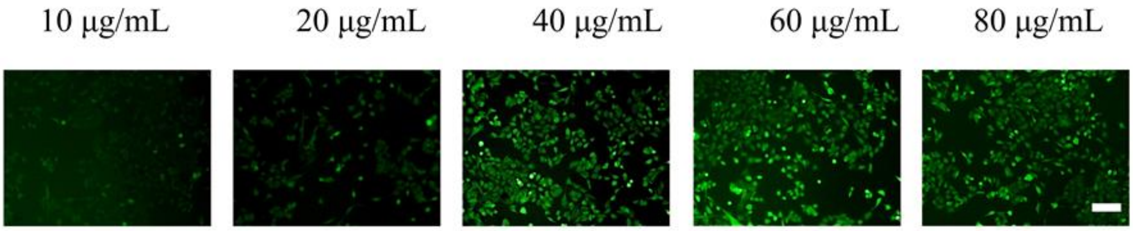

**Supplementary Fig. S24** ROS production was detected by DCFH-DA in Hepa1-6 cells at different concentrations. scale bar: 20 μm.

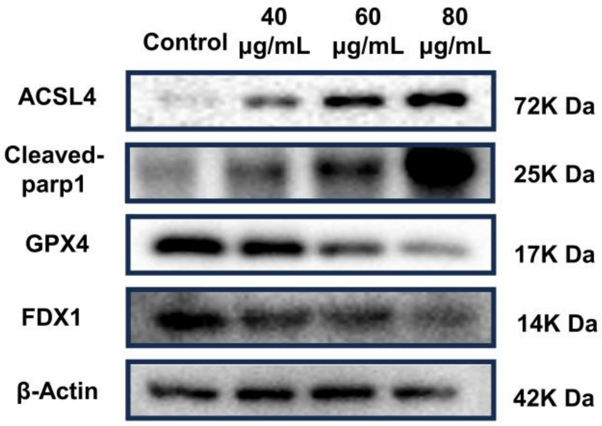

**Supplementary Fig. S25** Western blot analysis of GPX4, Cleaved- parp1, ACSL4, FDX1 expression after incubated with different concentrations of NMOF-Cu/Fe-Dox@M.

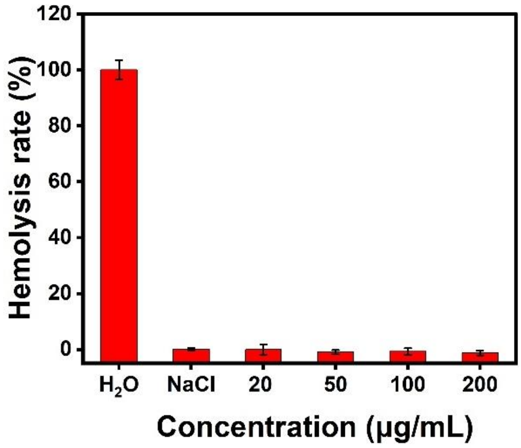

**Supplementary Fig. S26** The results of the hemolysis experiment of NMOF-Fe/Cu-Dox@M ($n$=3).

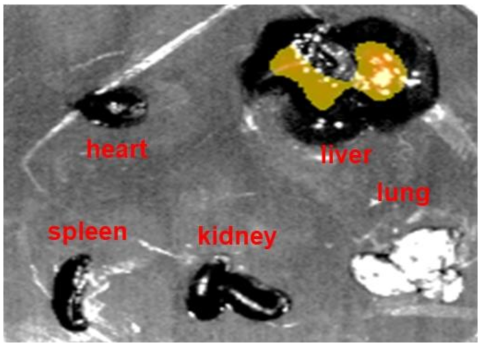

**Supplementary Fig. S27** Bioluminescence images of major organs of mice after tail vein injection of MOF-Fe/Cu-Dir@M for 12 h.

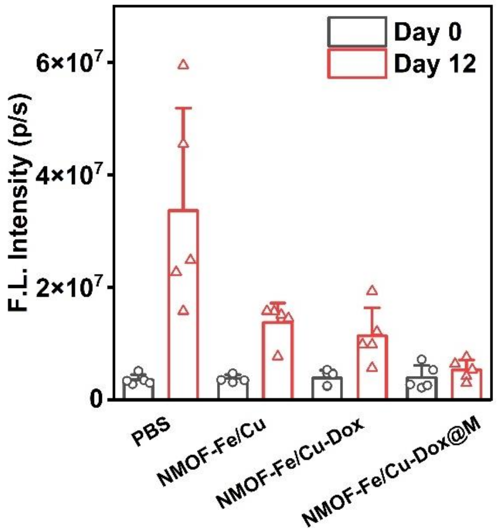

**Supplementary Fig. S28** The fluorescence intensity of various group of tumors in Day 0 and Day12 ($n$=5).

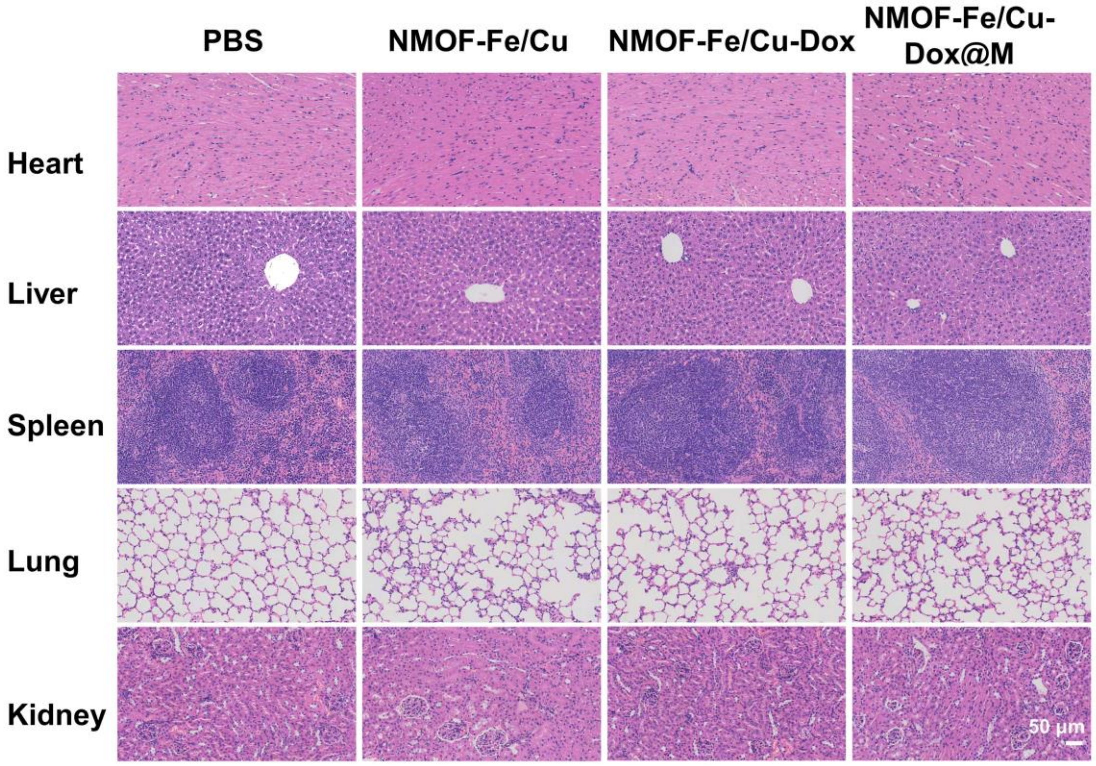

**Supplementary Fig. S29** H&E histopathological analysis of major organ tissues after various treatments.

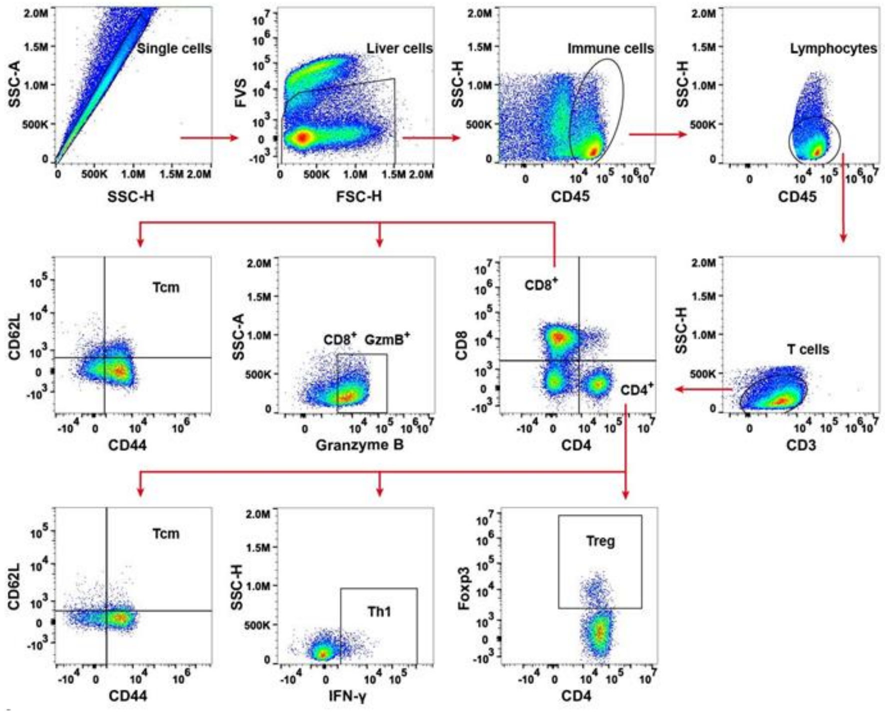

**Supplementary Fig. S30** Flow gating strategy of myeloid cells in the tumor.

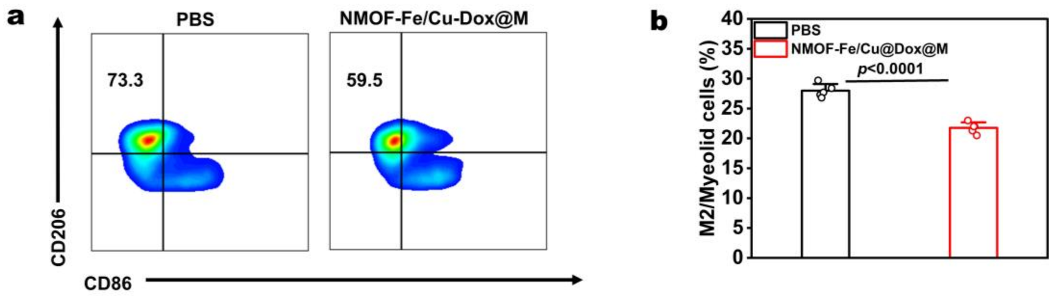

**Supplementary Fig. S31** Representative flow cytometric plots of M2 macrophages and quantification of M2 macrophages after different treatments ($n$=5). Statistical significance was calculated by $t$-test.

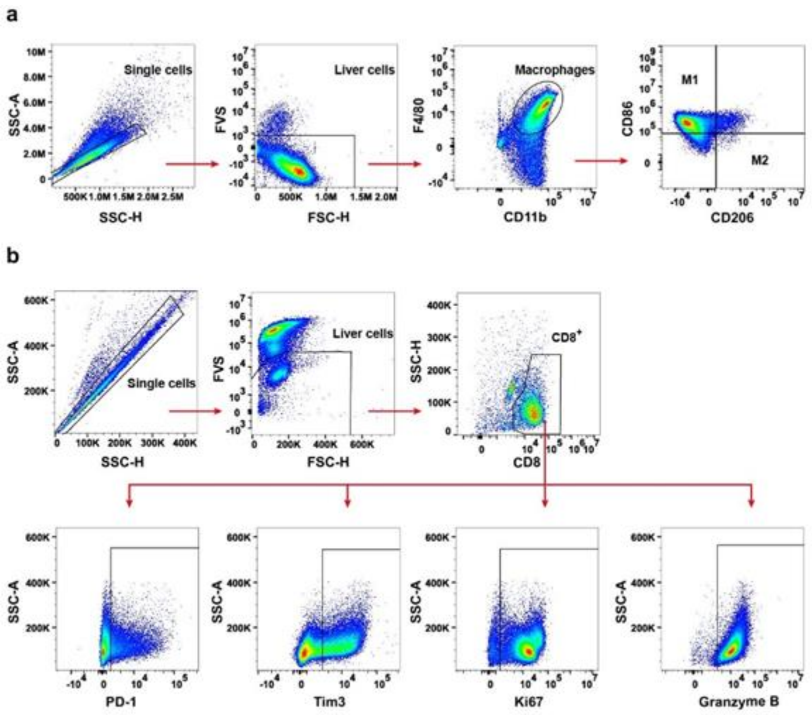

**Supplementary Fig. S32** Flow gating strategy of (a) macrophages and (b) $CD8^+$ T cells.

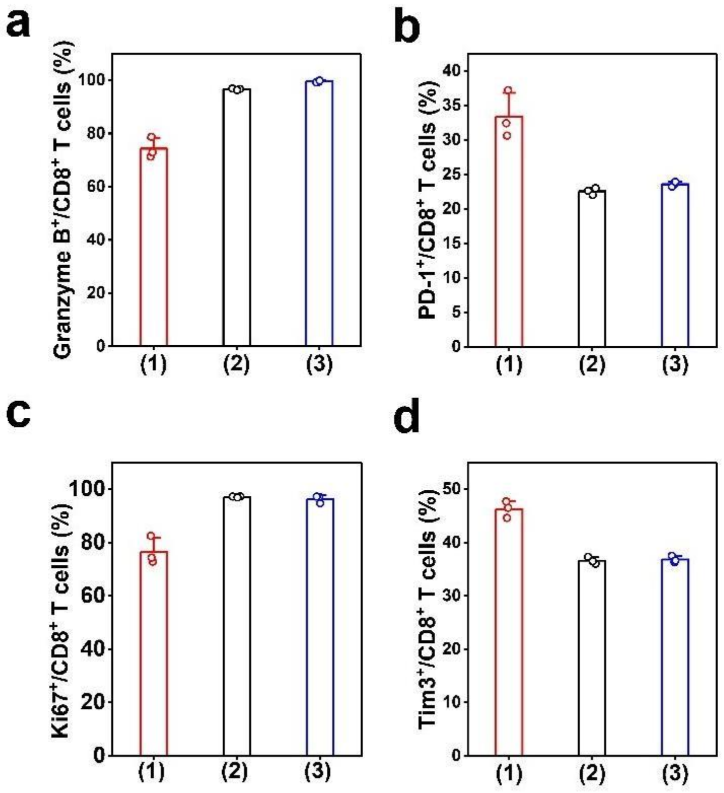

**Supplementary Fig. S33** Percentages of Granzyme $B^+$, and PD-$1^+$， $Ki67^+$ and $Tim3^+$ T cells in $CD8^+$ T cells extracted from the spleen cultured with RAW264.7 cells after co-culturing. (1) CM, (2) CM $_{\text{Hepa1-6}}$ and (3) CD3/CD28 -treated groups, respectively (*n*=3).

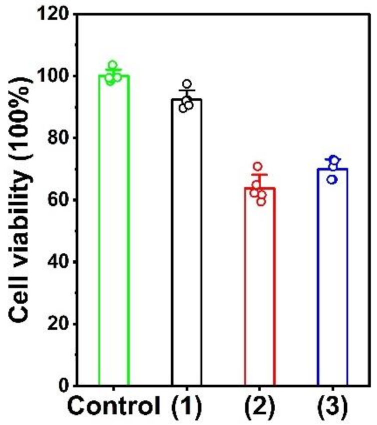

**Supplementary Fig. S34** The cytotoxic effects of co-cultured $CD8^+$ T cells on Hepa1-6 cells, (1) CM, (2) $CM_{Hepa1\text{-}6}$ and (3) CD3/CD28-treated groups, respectively (*n*=5).

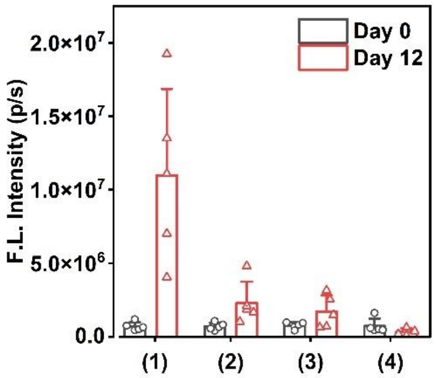

**Supplementary Fig. S35** The fluorescence intensity of various group of tumors in Day0 and Day12. (1) PBS; (2) NMOF-Fe/Cu-Dox@M; (3) αPD-1; (4) NMOF-Fe/Cu-Dox@M+αPD-1 (n=5).

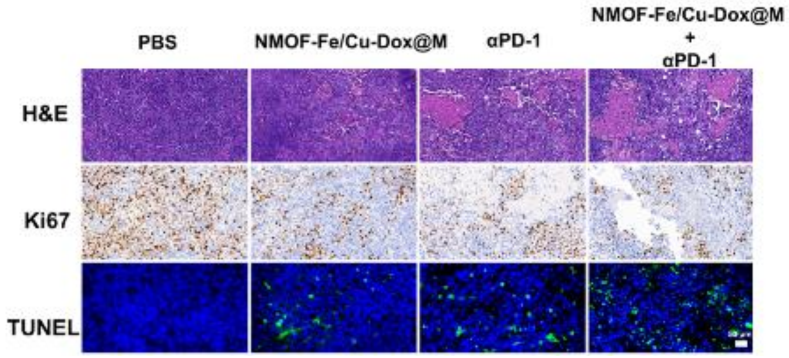

**Supplementary Fig. S36** Tumor tissue staining by H&E, Ki67, TUNEL after antitumor treatment and immunotherapy. Scale bar: 50 µm.

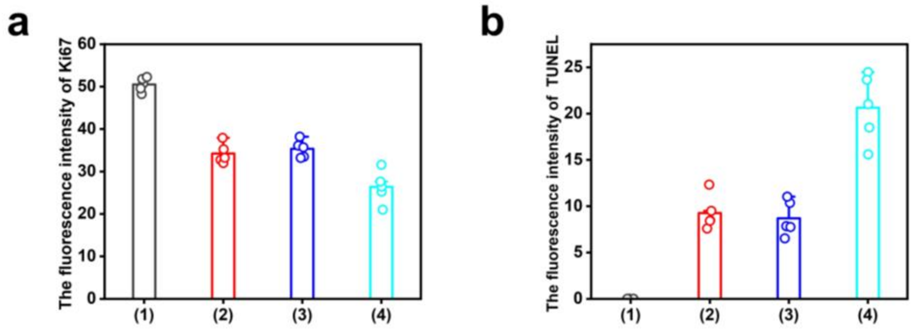

**Supplementary Fig. S37 (**a)The fluorescence intensity of Ki67 after various treatments. (b) The fluorescence intensity of Tunel after various treatments. (1) PBS; (2) NMOF-Fe/Cu-Dox@M; (3) αPD-1; (4) NMOF-Fe/Cu-Dox@M+αPD-1(*n*=5).

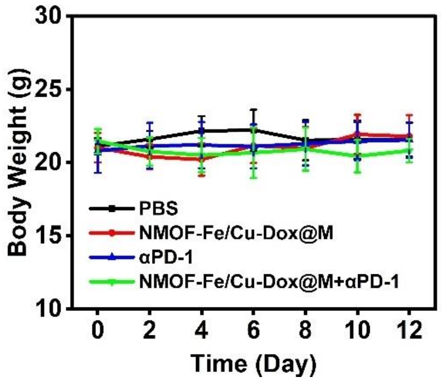

**Supplementary Fig. S38** The body weight variation in different treatments (n=5).

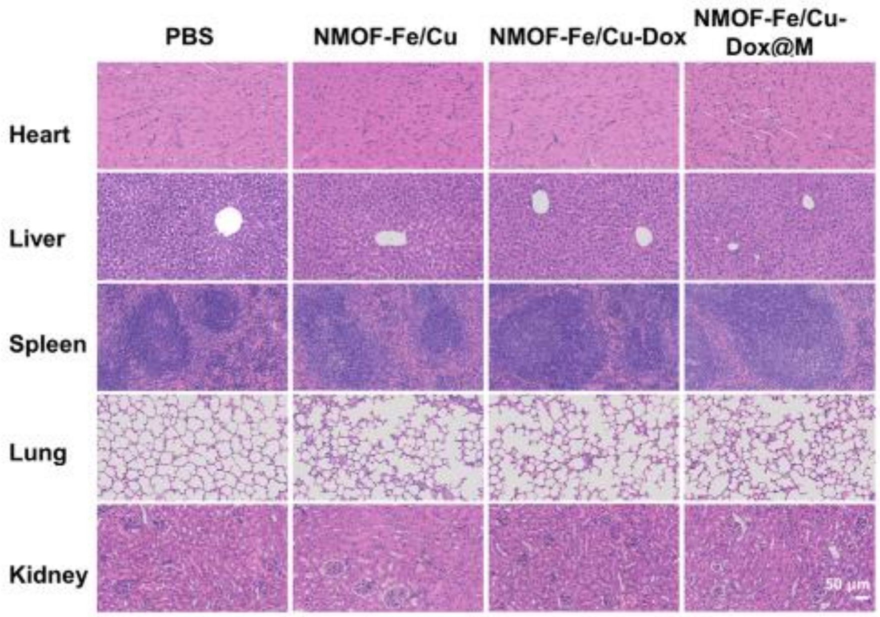

**Supplementary Fig. S39** H&E histopathological analysis of major organ tissues after various treatments. Scale bar: 50 μm.

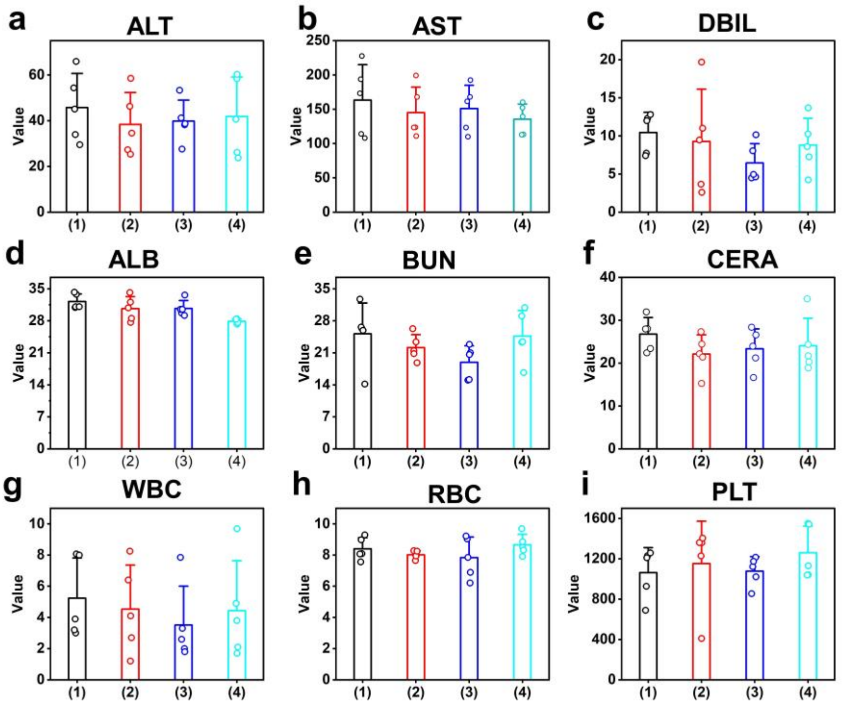

**Supplementary Fig. S40** The indices of liver function, renal function indicator and blood cell index in mouse serum after 12-day treatment. (1) PBS; (2) NMOF-Fe/Cu-Dox@M; (3) αPD-1; (4) NMOF-Fe/Cu-Dox@M+αPD-1 ($n$=5).

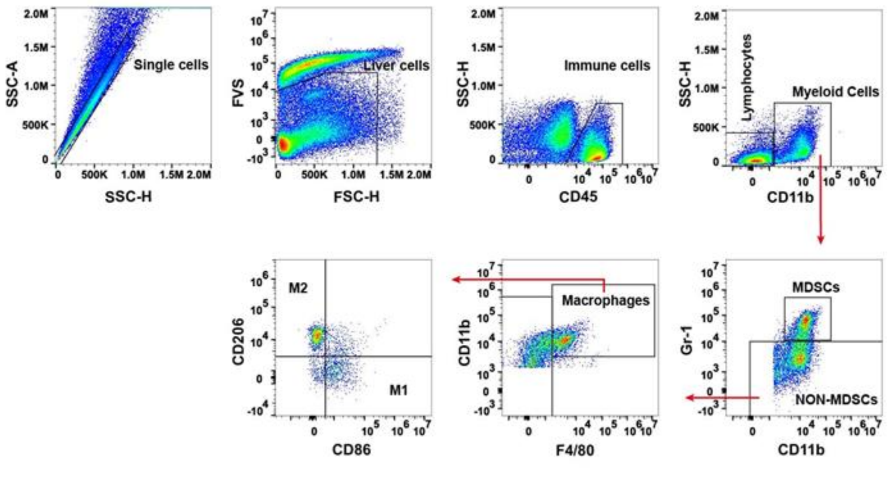

**Supplementary Fig. S41** Flow gating strategy of mouse immune T cells.

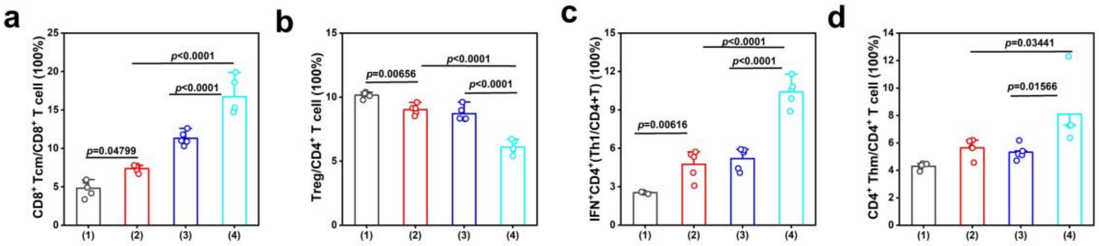

**Supplementary Fig. S42** The quantification of $CD8^+$ T cells (a), Treg cells (b), $CD4^+$/$IFN\gamma^+$ T cells (c), $CD4^+$ Tcm cells (d) surface marker expression in tumors of mice after different treatments (n=5). (1) PBS; (2) NMOF-Fe/Cu-Dox@M; (3) αPD-1; (4) NMOF-Fe/Cu-Dox@M+αPD-1 (*n*=5).

Supplementary table S1. The concentration of metal ion.

| metal ion | Liner equation | $R^2$ | μmol/mg |
|---|---|---|---|
| $Fe^{3+}$ | y=88316847.6844x | 0.9999 | 0.022 |
| $Cu^{2+}$ | y=214206900.3595x | 0.9989 | 1.21 |

Supplementary table S2. Antibodies and materials used for this work.

| *Antibody* | *Supplier* | N.O. |
|---|---|---|
| Ms CD8a FITC 53-6.7 | BD | 553030 |
| Ms IFN-Gma PerCP-Cy5.5 XMG1.2 | BD | 560660 |
| Ms CD62L PE-Cy7 MEL-14 | BD | 560516 |
| Ms Foxp3 Alexa 647 MF23 | BD | 560401 |
| Fixable Viability Stain 700 | BD | 564997 |
| Ms CD45 APC-Cy7 30-F11 | BD | 557659 |
| Ms CD3e BV421 145-2C11 | BD | 562600 |
| Ms CD44 BV510 IM7 | BD | 563114 |
| Ms CD4 BV605 RM4-5 | BD | 563151 |
| Ms Ly-6G/Ly-6C FITC RB6-8C5 | BD | 553126 |
| Ms NK-1.1 PE PK136 | BD | 553165 |
| PE-CF594 Rat Anti-Mouse CD19 | BD | 562291 |
| Ms Ly-6G PerCP-Cy5.5 1A8 | BD | 560602 |
| Ms CD11c PE-Cy7 HL3 | BD | 558079 |
| Ms CD206 Alexa 647 MR5D3 | BD | 565250 |
| CD11b BV510 M1/70 | BD | 562950 |
| Ms Ly-6C BV605 AL-21 | BD | 563011 |
| Ms F4/80 BV650 T45-2342 | BD | 743282 |
| BV786 Rat Anti-Mouse CD86 | BD | 740877 |
| Transcription Factor Buffer Set 100Tst | BD | 562574 |
| Leuko Act Cktl with GolgiPlug | BD | 550583 |
| Ms CD16/CD32 Pure 2.4G2 | BD | 553141 |
| PE-Cy™7 Mouse anti-Ki-67 | BD | 561283 |
| PE anti-human/mouse Granzyme B Recombinant Antibody | BioLegend | 372208 |
| Brilliant Violet 785™ anti-mouse CD279 (PD-1) Antibody | BioLegend | 135225 |
| Brilliant Violet 605™ anti-mouse CD366 (Tim-3) Antibody | BioLegend | 119721 |
| Mouse T-Activator CD3/CD28 | Gibco | 11452D |
| EasySep™ Buffer | STEMCELL | 20144 |
| EasySep™ Mouse CD8+ T Cell Isolation Kit | STEMCELL | 19853 |
| β-action | ABclonal | AC026 |
| ACSL4 | ABclonal | A20414 |
| Cleaved-parp1 | ABclonal | A19612 |
| GPX4 | ABclonal | A11243 |
| DLAT | proteintech | 68303-1-Ig |
| FDX1 | Zen-bioscience | R23394 |
| PVDF membrane-0.22μm | Merck | IPVH00010 |
| PVDF membrane-0.45μm | Merck | ISEQ00010 |

| | | |
|---|---|---|
| Ki67 | Servicebio | GB121141 |
| CD206 | Cell Signaling Technology (CST) | 24595 |
| CD8 | Servicebio | GB1506 |
| GranzymeB | Cell Signaling Technology (CST) | 44153 |
| F4/80 | Cell Signaling Technology (CST) | 70076 |
